# Evidencing the role of a conserved polar signaling channel in the activation mechanism of the μ-opioid receptor

**DOI:** 10.1101/2024.08.20.608751

**Authors:** Arijit Sarkar, Szabolcs Dvorácskó, Zoltán Lipinszki, Argha Mitra, Mária Harmati, Krisztina Buzás, Attila Borics

## Abstract

The activity of G protein-coupled receptors has been generally linked to dynamically interconverting structural and functional states and the process of activation was proposed to be controlled by an interconnecting network of conformational switches in the transmembrane domain. However, it is yet to be uncovered how ligands with different extent of functional effect exert their actions. According to our recent hypothesis, the transmission of the external stimulus is accompanied by the shift of macroscopic polarization in the transmembrane domain, furnished by concerted movements of conserved polar amino acids and the rearrangement of polar species. Previously, we have examined the μ-opioid, β_2_-adrenergic and type 1 cannabinoid receptors by performing molecular dynamics simulations. Results revealed correlated dynamics of a polar signaling channel connecting the orthosteric binding pocket and the intracellular G protein-binding surface in all three class A receptors. In the present study, the interplay of this polar signaling channel in the activation mechanism was evidenced by systematic mutation of the channel residues of the μ-opioid receptor. Mutant receptors were analyzed utilizing molecular dynamics simulations and characterized *in vitro* by means of radioligand receptor binding and G protein stimulation assays. Apart from one exception, all mutants failed to bind the endogenous agonist endomorphin-2 and to stimulate the G_i_ protein complex. Furthermore, mutation results confirmed strong allosteric coupling between the binding pocket and the intracellular surface. The strong association and optimal bioactive orientation of the bound agonist was found to be crucial for the initiation of correlated motions and consequent signaling.

## 1. Introduction

G protein-coupled receptors (GPCRs) are one of the most coveted categories of drug targets for therapeutic interventions, evidenced by the fact that roughly one-third of all prescription drugs aim to affect receptors within this family. [1,2] The use of certain GPCR-targeting drugs is, however, limited by undesired side effects which might result from non-selective activation of multiple GPCRs or multiple signaling pathways associated with a single receptor. A current challenge is to develop functionally selective drugs, specific to a particular signaling pathway, albeit. This is complicated by the fact that the structural mechanism of activation of these receptors as well as the ligand-receptor dynamics is still largely unclear. Only a few varieties of G proteins are involved in signaling, with the presence of ligands in far more excess in comparison, which suggests that GPCR activation may follow a general mechanism. This information is critical in designing drugs to selectively target specific signaling pathways, thereby reducing side effects and improving therapeutic outcomes. According to the common theory of GPCR activation, the receptor undergoes conformational changes upon binding to agonists that activate downstream signaling pathways. These pathways include the G protein-coupled pathway, which involves the activation of G proteins and subsequent regulation of secondary messengers such as cyclic adenosine monophosphate (cAMP) [3] as well as other intracellular signaling pathways, such as the mitogen-activated protein kinase (MAPK) [4] and phospholipase C (PLC) pathways. [5]

Structurally, GPCRs are transmembrane proteins which consist of a conserved domain of seven transmembrane helices (TM domain) connected by intra and extracellular loops, an extracellular N-terminal domain, and an intracellular C-terminal domain. In the recent years, there has been significant research interest in unraveling the structure of GPCRs in high spatial resolution, leading to the advancement of cutting-edge techniques. From the traditional x-ray and lipidic cubic phase (LCP) crystallography to emerging methodologies such as single-particle cryo-electron microscopy (cryo-EM) and x-ray free-electron laser (XFEL) protein crystallography, a plethora of state-of-the-art techniques have been developed for this purpose. Such developments have enabled researchers to obtain high-resolution structures of GPCRs. [6] The knowledge of the process of transitioning from the inactive state to the active signaling state of GPCRs has become increasingly complex as novel discoveries continue to emerge each year. Upon activation of GPCRs, there is a remarkable level of structural conservation in the intracellular changes that occur, indicating a common evolutionary origin for the activation mechanism across most GPCRs. The above studies as well as landmark molecular dynamics simulation results indicated that the activation process of class A GPCRs is accompanied by the displacement of the 6^th^ transmembrane helix (TM6), resulting in the creation of a cavity on the receptor’s intracellular face that can accommodate the C-terminus of the G protein α-subunit. Additionally, transmembrane helix 5 (TM5) also rotates away from the receptor, further enlarging the cavity for G protein binding. The dynamic nature of such receptors can be showcased by the suggestion that GPCRs can exist in dynamically interconverting active, inactive and intermediate structural states, even in the absence of ligands. The population of such states relies upon ligand binding and orientation, physico-chemical properties of the ligand, interacting intracellular signaling proteins and lipid membrane composition, among other factors. [7–12] Recent studies have shed light on the potential role of several other variables on activation. Key conserved functional motifs, including E/DRY, NPxxY, and CWxP, are recognized to play a role in the activation process of class A GPCRs. Additionally, when TM helices undergo reorganization, it is usually accompanied by the synchronized interplay between the mentioned conserved motifs, and a network of water molecules present in the inner cavities of the TM domain. The continuous polar pathway, created by this network, functions as a linkage between the ligand binding pocket and the intracellular G protein binding surface. It has been suggested as the fundamental pathway of receptor activation. Furthermore, the elevated levels of Na^+^ leading to inhibition of agonist-induced activation of opioid receptors and related GPCRs, due to a preserved allosteric Na^+^ binding site was revealed. [13–20] Earlier studies have provided evidence that the receptors adapt ’canonical active’ and ’alternative active’ conformations, linked to G protein-mediated signaling and β-arrestin-mediated signaling respectively, contributing to the theory of biased signaling. [21,22] Typically, distinct active states mentioned are characterized by specific pairwise residue-residue distances and side-chain conformations. These changes often entail a counter-clockwise twist at the 7^th^ transmembrane helix (TM7) and a shift in the allosteric pocket related hydrogen bonding network. This evidence is a major step forward toward biased ligand design, which could alleviate undesired side effects. However, explanations that rely solely on a structural basis may not offer a clear and definitive resolution for the mechanism of GPCR activation, especially in situations where ligands with similar structural features and affinities, but distinct functional properties and efficacies are involved.

The μ-opioid receptor (MOP) is a class A GPCR that initiates intracellular signaling cascades upon binding to endogenous or exogenous opioids. These signaling pathways are critical for the analgesic effects, but also contribute to the unwanted side effects of exogenous opioids, such as development of tolerance, dependence, and addiction [23,24] Several x-ray crystallographic and cryo-EM structures of MOP have been determined, revealing important insights into the receptor’s structure and function. [25–32] These structures have shed light on how the receptor interacts with a variety of ligands and how these interactions result in the activation of the receptor, providing a basis for the design of new drugs that could target MOP with greater specificity and fewer side effects. Overall, the study of MOP and its experimental structures has important implications for our understanding of pain management and addiction, as well as for the development of new drugs to overcome the ‘opioid crisis’ faced by our society.

Results of a recent, extensive MD simulation study involving the MOP in active and inactive states showed highly correlated internal motions among the side chains of the conserved functional motifs; the orthosteric and allosteric binding pockets, the NPxxY motif, and the cytosolic helix (H8), mentioned earlier, which form a polar signaling channel connecting the ligand binding pocket to the intracellular surface. These ordered synchronous motions were exclusively observed in the active state MOP when bound to the G_i_ protein and not in the inactive state, or when bound to the β-arrestin-2 complex, indicating that these motions could be associated with G protein-mediated signaling. [33] Follow-up MD simulation studies involving two different class A GPCRs corroborated the above findings, suggesting the existence of a generalized activation pathway for class A GPCRs. [34,35] To address the uncertainties around biased signaling an auxiliary hypothesis was formulated that states, that receptor activation is accompanied by a shift of electrostatic balance in a central duct of the TM domain. The electrostatic balance in the orthosteric pocket is perturbed upon ligand binding which perturbation is then propagated to the intracellular surface of the receptor through the synchronous movements of conserved polar amino acid side chains, assisted by water molecules, leading to G protein dissociation and subsequent downstream signaling. The hypothesis could be supported by the fact that mutations affecting the polarity of functional motifs in class A GPCRs can have a significant impact on their function as well as their ligand binding affinity. [17,28,36–45] Additionally, Na^+^ binding in the allosteric site has been shown to inhibit the activation of many class A GPCRs, further suggesting the importance of charge balance and polarity changes in receptor signaling. Mutation studies have revealed that the residues which coordinate Na^+^ play a crucial role in determining the efficacy of a drug, and therefore can be a promising target for drug development. [17,19]

In this research, we investigated the mechanisms underlying MOP activation and observed the effect of altering the polarity of the proposed signaling channel on ligand binding and the signaling activity of the receptor. Multiple mutants of the MOP receptor were designed and generated, in which the polarity of signaling channel constituents was modified to different extents. The effects of mutations were examined via simulation and experimental approaches.

## 2. Methods

### 2.1 Design of mutant MOP receptors

Following up on our study involving MOP [32], point mutations were introduced in the participants of the polar signaling channel, namely Y326^7.43^F, N328^7.45^D, N328^7.45^L, D340^8.47^N, and D340^8.47^L. These residues belong to the orthosteric pocket, the allosteric Na^+^ anchoring site, and the intracellular G protein coupling interface, respectively (Figure 1). In previous studies involving different GPCRs, mutations of the elements of the polar signaling channel led to impaired G protein signaling or increased constitutional activity. [17,36–45] The choice of the mutants for our investigation was based on their positions, and lack of extensive studies for most of the residues concerning MOP.

**Figure 1.**
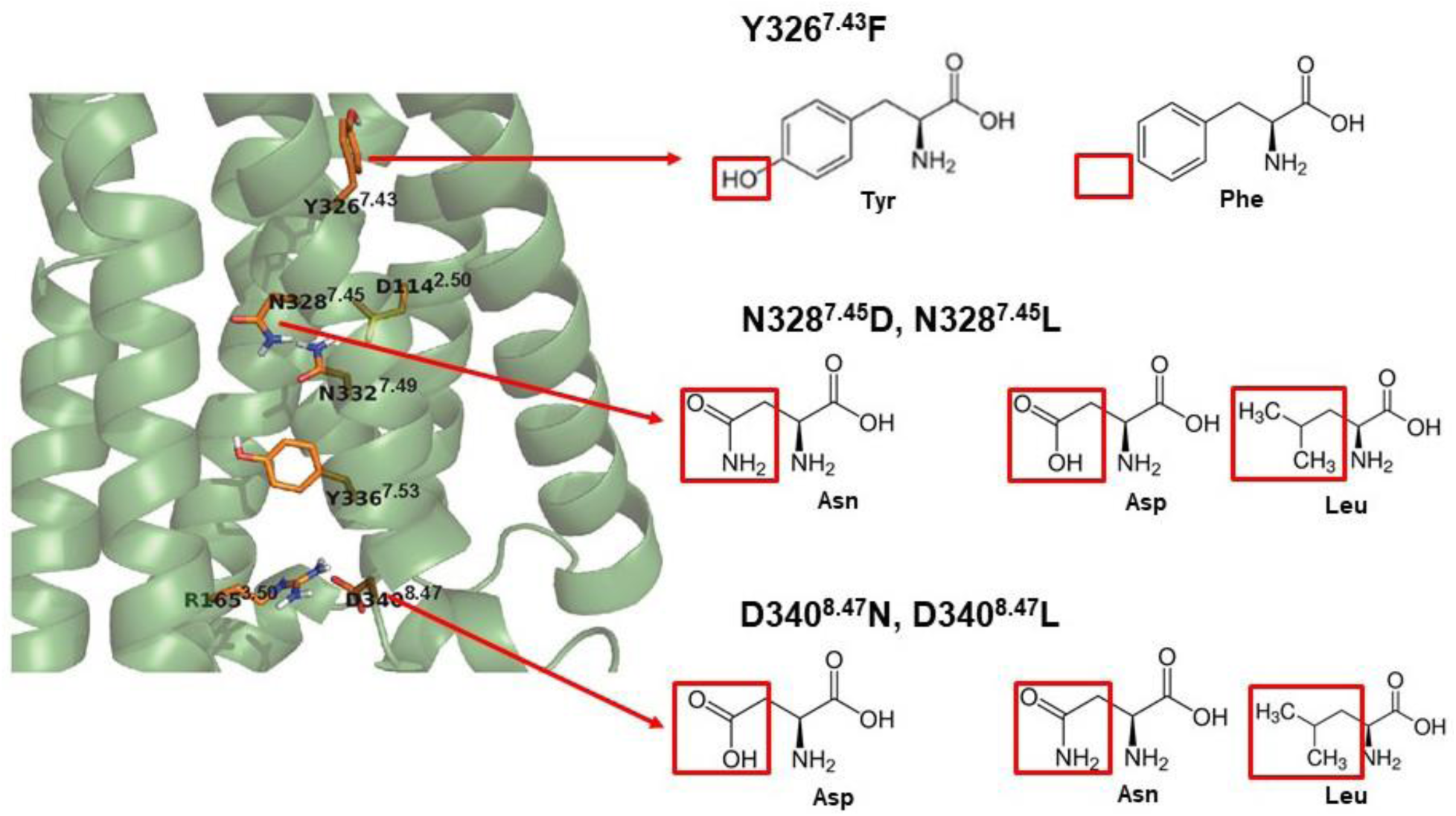
Selected mutations aimed at altering the polarity of the proposed signaling channel of the MOP.

### 2.2 Preparation of simulation systems

The cryo-EM structure of the active state MOP-G_i_ protein complex (PDB code: 6DDE), [10] was obtained from the Brookhaven Protein Data Bank (http://www.rcsb.org). Additionally, separate coordinates of the G_i_ protein heterotrimeric complex and GDP (PDB code: 1GP2) [46] were used as essential components in the simulations. GTP was generated in CHARMM-GUI [47] and replaced GDP from the G_i_ complex. The crystallographic structures of the G_s_ protein-bound active β_2_-adrenergic (β_2_AR, PDB code: 3SN6) [48] were utilized as a template to orient the G_i_ protein complex to the receptor. The crystallization chaperones and fusion proteins were excluded from the starting structures. Single point mutants of the MOP were generated virtually, using the mutagenesis function of Pymol 2.4.0., keeping focus on the effect of alteration of the polarity of the established signaling channel. Specifically, Y326^7.43^ in the ligand binding pocket was mutated to F; N328^7.45^, a component of the allosteric Na^+^ binding pocket, was mutated to D and L, and D340^8.47^, a part of the intracellular G protein coupling interface, was mutated to N and L (Fig. 1).

The sequences of the murine MOP (UniProtKB-P42866-OPRM1) were retrieved from the UniProt database (http://www.uniprot.org). 10 ns folding simulations were performed to reconstruct the flexible N-and C-terminal domains which are missing from their experimental structures. The simulation utilized the GROMACS 2018.3 program package, [49] the AMBER ff99SB-ILDN-NMR force field, [50] and the GB/SA implicit solvation model, [51] in accordance with a previously established protocol. [33] Any additional missing, modified, or mutated residues within the structure of the receptor were restored or corrected using the Swiss-PdbViewer program ver. 4.10. [52] Throughout MD simulations, the system temperature was set to 310 K, and v-rescale algorithm was used for its regulation. [53] Ten parallel simulations were conducted for both the terminal domains, and the resulting folded structures were assessed and chosen based on their compactness and the accessibility of post-translational modification (PTM) and TM region attachment sites. The suitably folded terminal domains were then appended manually to the TM region using Pymol 2.4.0.

The PTMs in the receptor structures, including glycosylation of the N-terminal domain, were incorporated using the CHARMM-GUI web-based platform. [47] The glycosylation involved a common core structure (Manα1–3 (Manα1–6) Manβ1–4GlcNAcβ1–4GlcNAcβ1–N) along with sialic acid (N-acetylneuraminic acid). Phosphorylation of serine and threonine residues in the C-terminal domain, along with palmitoylation of a cysteine residue in the 3^rd^ intracellular loop (ICL3), were also introduced. A single disulfide bridge between two cysteine residues, which play a critical role in stabilizing the structure of the protein was also added. The sites of the PTMs are noted in Table S1.

The structure of endomorphin-2 (EM2), an endogenous peptide agonist of the MOP [53] was constructed manually using Pymol 2.4.0. Since these simulation systems were built prior to the publication of the cryo-EM structure of MOP bound endomorphin-1 (EM1), a peptide agonist analogous to EM2, [28,55] molecular docking was utilized to predict the binding site and orientation of EM2 in MOP. The Autodock 4.2 software, [55] employing the Lamarckian genetic algorithm, was used for the flexible docking of EM2 to the active state MOP experimental structure, to verify its orthosteric binding site. The docking was conducted within an 8.0 nm × 8.0 nm × 8.0 nm grid volume, ensuring comprehensive coverage of the receptor surface accessible from the extracellular side, including the ligand binding pocket. All φ, ψ, and χ^1^ ligand torsions, as well as receptor side chains in contact with the bound ligand (D147^3.32^, Y148^3.33^, M151^3.36^, K233^5.39^, W293^6.48^, I296^6.51^, H297^6.52^, V300^6.55^, I322^7.39^, Y326^7.43^), were kept flexible during the docking procedure. The grid points were spaced at intervals of 0.0375 nm, and a total of 1000 dockings were performed. The resulting ligand-receptor complexes underwent clustering and ranking based on their corresponding binding free energies. The lowest energy bound state, indicating the presence of the specific ligand-receptor interactions observed in the cryo-EM structure of MOP with peptide agonist DAMGO, was selected verifying the optimal localization of EM2 in the pocket. The orientation of EM2 was meticulously scrutinized to adhere to its bioactive conformation, drawing insights from both the experimental structure and our studies. [10,32,33] The cryo-EM structure of the MOP - EM1 complex published later have validated the orientation of EM2 used in our studies.

To emulate their native environment, membrane bilayers with caveolar compositions were constructed based on lipidomic data, [56] also validated by our earlier studies. [33–35] The membrane builder tool of CHARMM-GUI was used to build the membrane system with the lipid components being parameterized according to CHARMM36m parameters. [47] The receptor complexes were incorporated into a resultant asymmetric multicomponent, raft-like membrane systems composed of cholesterol (CHL∼32.8%), 1-palmitoyl-2-oleoyl-glycero-3-phosphatidylcholine (POPC∼14.9%), 1-palmitoyl-2-oleoyl-sn-glycero-3-phosphatidylethanolamine (POPE∼27.8%), 1-palmitoyl-2-oleoyl-sn-glycero-3-phosphatidyl-L-serine (POPS∼3.6%), 1-palmitoyl-2-oleoyl-sn-glycero-3-phosphatidylinositol (POPI2∼6%), palmitoyl-sphingomyelin (PSM∼9.9%) and monosialodihexosylganglioside (GM3∼5%). According to recent literature data, the asymmetric composition of the lower and upper leaflets was determined. [57] The systems were explicitly solvated with TIP3P water molecules in a hexagonal-shaped periodic box. To achieve physiological ionic strength, Na^+^ and Cl^-^ ions (0.15 mM) were added and the net charge of the system was set to neutral. The coordinates and topologies of the systems were generated in GROMACS format, with CHARMM36m all-atom force field parameters assigned to the generated components. [58] Altogether, a total of 12 independent 1 µs simulations were set up in the workflow for the MOP complexes.

### 2.3 MD simulations

Following the building of a caveolar membrane around the receptor complexes, the GROMACS 2018.3 program package [49] was utilized to perform thorough energy minimization and subsequent equilibration of the whole system, closely following the protocol established in our previous works. [33–35] The simulation systems were subjected to an initial 5000 steps of steepest descent, and subsequent 5000 steps of conjugate gradient minimization. The convergence criterion for both steps was set at 1000 kJ mol^-1^ nm^-1^. Using a six-step protocol supplied by CHARMM-GUI, the equilibration process involved two consecutive MD simulations at 303.15 K in the canonical (NVT) ensemble, followed by four additional simulations in the isobaric-isothermal (NPT) conditions at the same temperature and 1 bar pressure. Positional restraints were imposed on the heavy atoms of the proteins and membrane constituents and gradually decreased throughout the equilibration protocol. The initial three equilibration MD runs were 25 ps in duration and conducted with 1 fs time steps. The subsequent two runs continued for 100 ps with 2 fs time steps, and the final equilibration step was extended to 50 ns and was executed in 2 fs time steps, similar to the approach described previously. [33–35] The correct bond lengths were maintained using the LINCS algorithm, temperature was regulated using the v-rescale algorithm [53] with a coupling constant of 1 ps, and pressure control was achieved using the c-rescale barostat. [59] The pressure coupling method had a coupling constant of 5 ps and isothermal compressibility of 4.5 × 10^−5^ bar^−1^. Energy contributions from long-range electrostatic interactions were calculated using the particle mesh Ewald (PME) summation method, and van der Waals interactions were computed with a twin-range cutoff. 1.2 nm was set as the cut-off value, as described earlier. [33–35]

Executing GROMACS 2018.3, unrestrained production simulations for the wild type receptor and its mutant systems were carried out in the NPT ensemble at 310 K, employing parameters akin to those mentioned above. Each production simulation extended for 1 µs. System coordinates were recorded every 5000 steps, yielding 100,000 snapshots per trajectory. Simulations of the active state wild type receptor as control were performed as well as simulations of its mutants mentioned earlier, complexed with the heterotrimeric G_i_ protein. These simulations were performed in the presence of the endogenous agonist EM2, oriented in its bioactive conformation (see above). The simulations were conducted in 2 sets, with the ligand subjected to mild positional restraints in the binding pocket (200 kJ mol^−1^ on heavy atoms) in one set of simulations and the ligand being allowed to be free without any restraints in the other. This approach aimed to prevent spontaneous ligand ejection from the pocket and assess the influence of native dynamics of the ligand, over it being in its static state, even in its functional form.

### 2.4 MD trajectory analysis

The analysis suite of the GROMACS 2018.3 package was used for the analysis of our output MD trajectories as outlined in our previous studies. [33–35] The radius of gyration of terminal domains, along with the measurement of the minimum distance between them was performed, to test the integrity of the simulation system. Potential artifacts caused by the proximity of the flexible terminal domains and their periodic replicas were also evaluated via these methods. The minimum distance between Na^+^ from the allosteric site was also assessed to report its penetration and localization. Root mean square deviation (RMSD) of the protein backbone atoms was measured to investigate the structural deviation of domains with respect to the starting experimental structures. Specific interactions between conserved microswitches were calculated by analyzing the presence of H-bonds as well as salt bridges between the residues. For the H-bond, the donor-acceptor distance and angle cutoff were set to 0.35 nm and 30.0 degrees or below, respectively; whereas for the salt bridge calculations, the distance and angle cut-off between the acidic and basic moieties were set to 0.4 nm and 90.0 degrees. The secondary structures of receptor domains were assigned and monitored during simulations using the the DSSP (Define Secondary Structure of Proteins) method. [60] Side chain rotamer conformations were also measured to determine the signaling state of the receptor. The correlation of side chain motions was evaluated using generalized cross-correlation matrix (GCCM) analyses based on linear mutual information (MI), using the g_correlation utility of the GROMACS 3.3 program package, following the methodology used in our previous work. [33–35] Briefly, correlation matrices were converted to heat maps using the GIMP ver. 2.10.30 software, with red intensity levels corresponding to MI values greater than 0.63. Visualization of molecular structures was carried out using Pymol 2.4.0, and graphs were created using the Xmgrace ver 5.1.25 program.

### 2.5 DNA constructs

The full-length complementary DNA (cDNA) of the mouse MOP receptor (GenBank #: AB047546.1), which includes the coding sequence (CDS) as well as the 5’ and 3’ untranslated regions (UTRs), was amplified using polymerase chain reaction (PCR) with Phusion Plus DNA polymerase (Thermo Fisher Scientific, Waltham, MA, USA, #F630). Genomic DNA extracted from the CHO-MOP cell line [61] (kindly provided by András Váradi, Memorial Sloan-Kettering Cancer Center, New York, United States) served as the template. The PCR product was blunt-end cloned into the pJET2.1 plasmid according to the manufacturer’s instructions (Thermo Fisher Scientific, #K1231). Verification of the wild-type MOP (MOP WT) was confirmed by DNA sequencing. Modified MOP constructs, each with a specific amino acid substitution (Y326^7.43^F, N328^7.45^D, N328^7.45^L, D340^8.47^N, or D340^8.47^L), were generated using the QuickChange II XL Site-Directed Mutagenesis Kit (Agilent Technologies, Santa Clara, CA, USA, # 200523). The cDNAs of these MOP variants were then PCR-amplified from the pJET2.1 plasmids and inserted into the BamHI site of the pTRE2hyg mammalian expression vector (Clontech, Palo Alto, CA, USA, #6255-1) using In-Fusion cloning (Takara Bio, Shiga, Japan, #639648). All constructs were confirmed by DNA sequencing before transfection. The oligonucleotide primers used are listed in Table S2.

### 2.6 Generation of stably transfected CHO-K1 Tet-On Cell Lines

To generate stable CHO-K1 Tet-On cell lines expressing MOP derivatives, purified plasmid DNA containing either WT or mutated forms of MOP cDNA in the pTRE2hyg vector was transfected into CHO-K1 Tet-On cells (Clontech, #C3021-1) using Torpedo-DNA transfection reagent (Ibidi, Gräfelfing, Germany, #60611) following the manufacturer’s guidelines, as previously described [62]. An empty pTRE2hyg vector was used to establish the control cell line. Selection of stably transfected cell lines began 24 hours post-transfection in low glucose-containing DMEM medium (Biosera, Nuaille, France, #LM-D1102), supplemented with 10% tetracycline-free fetal bovine serum (Biosera, #FB-1001T), 1 × Penicillin-Streptomycin (Biosera, #XC-A4122), 2 mM stable glutamine (Biosera XC-T1755), 100 µg/ml G418 sulfate (Corning, NY, USA, #61-234), and 250 µg/mL Hygromycin B (Corning, #30-240-CR). The selection process continued for 2–3 weeks with regular medium changes until distinct colonies were visible. Cells were cultured in 6 cm or 10 cm tissue culture dishes in a 37 °C humidified incubator with 5% CO_2_. Transgene expression was induced by treating the cells with 1.5 µg/mL doxycycline (Clontech, cat# 631311) for 24 hours.

### 2.7 Membrane preparation

Membranes of CHO-K1 cells stably expressing the wild type and mutant MOP were prepared from 4 monolayer cell cultures in 175 cm^2^ flasks. Cells were rinsed with DPBS 3 times and dissociated with 0.5 mM EDTA (Merck Millipore, Burlington, MA, USA, #324503) for 8-10 min at room temperature. Then, the cells were washed with DPBS (Lonza, # 17-515F) via centrifugation at 1500 × g for 10 min at 4 °C. Cell pellet was resuspended in 50 mM Tris–HCl (pH 7.4), frozen at -80 °C for 1 h and homogenized with Dounce homogenizer after thawing. Homogenate was ultracentrifuged at 100,000 × g for 30 min at 4 °C. Then, the pellet was resuspended in 50 mM Tris–HCl (pH 7.4) and stored in aliquots at -80 °C. Protein concentration of the membrane preparations was measured with the Pierce™ BCA Protein Assay Kit (Thermo Fisher Scientific, #23225) based on the manufacturer’s instructions.

### 2.8 Radioligand competition binding assay

Binding assays were carried out following a previously published protocol [63], at 25 °C for a duration of 60 minutes using a 50 mM Tris–HCl buffer (pH 7.4) in glass tubes. Each assay had a total volume of 1 mL and contained 0.5 mg/mL of membrane protein. For competition binding assays, cell membranes were exposed to 2 nM of [^3^H]DAMGO (K_d_: 0.5 nM) along with increasing concentrations (10^−12^ - 10^−5^ M) of synthetic, unlabeled EM2 (generously provided by Géza Tóth from the Institute of Biochemistry, HUN-REN Biological Research Centre, Szeged, Hungary) [64]. Non-specific binding was evaluated using 10 μM naloxone. Following incubation, samples were diluted with an ice-cold wash buffer (50 mM Tris–HCl, pH 7.4) and then subjected to multiple washes. The samples were rapidly filtered through Whatman GF/C glass fiber filters (Whatman Ltd., Maidstone, UK) using a 24-well Brandel Cell Harvester (Gaithersburg, MD, USA). The filters were subsequently air-dried, immersed in Ultima Gold MV scintillation cocktail, and the radioactivity was quantified with a TRI-CARB 2100TR liquid scintillation analyzer (Packard, Perkin Elmer, Waltham, MA, USA).

### 2.9 Ligand stimulated [^35^S]GTPγS binding assay

The ligand-stimulated [^35^S]GTPγS binding assays were performed as outlined previously [63]. Briefly, cell membranes (30 μg protein/tube) were prepared and incubated with 0.05 nM of [^35^S]GTPγS (PerkinElmer) and varying concentrations (10^−11^ - 10^−5^ M) of unlabeled EM2 in a buffer containing 30 μM GDP, 100 mM NaCl, 3 mM MgCl_2_, and 1 mM EGTA in 50 mM Tris– HCl buffer (pH 7.4). The incubation was performed at 30 °C for 60 minutes. Basal [^35^S]GTPγS binding was recorded in the absence of ligand and was used as a reference value (100%). Non-specific binding was determined by the addition of 10 μM unlabeled GTPγS, which was then subtracted from the total binding. The processes of incubation, filtration, and measurement of radioactivity were executed as described above.

### 2.10 Data analysis

The calculation of inhibitory constants (K_i_) during competition binding assays was conducted by analyzing the inflection points of the displacement curves through nonlinear least-squares curve fitting, as previously reported. [65] These constants were determined using the Cheng-Prusoff equation: K_i_ = EC_50_/(1 + [ligand]/K_d_). For the [^35^S]GTPγS binding assays, data were reported as the percentage increase in specific [^35^S]GTPγS binding compared to basal levels. Each assay was performed in triplicate, and potency (EC_50_) and efficacy (E_max_) were assessed using sigmoid dose-response curve fitting methods, consistent with the procedures detailed in our earlier publication. [65] Statistical analysis of E_max_ and EC_50_ values was carried out using one-way ANOVA followed by Tukey’s multiple comparison test (****P < 0.0001) and unpaired Student’s t-test. Furthermore, comparisons of [^35^S]GTPγS binding values for EM2, in the presence or absence of 1 and 10 μM naloxone, were performed using one-way ANOVA with subsequent Tukey’s test (****P < 0.0001, **P < 0.01, *P < 0.05). All data analyses and curve fitting were executed using GraphPad Prism 9.0 Software (San Diego, CA, USA), as described earlier. [65]

## 3. Results and discussion

### 3.1 Simulation system integrity

Building upon the methodology employed in our previous studies [33–35], we adopted a similar approach in constructing the simulation systems. Particular attention was given to incorporating both the N- and C-terminal domains of the receptor while constructing the simulation systems, to achieve a realistic approximation of the native structure. This inclusion was crucial for accounting for the drag caused by the mass of these domains and its subsequent impact on the dynamics of the transmembrane helices whose significance in the activation of the receptor has been greatly emphasized. [7,15,20] Consequently, their accurate representation was deemed essential in the simulation systems developed for this study.

During the course of simulations, a notable observation was the frequent occurrence of partial unfolding in the N- and C-terminal domains, evidenced by the changes in the radii of gyration over time (Figure S1). It was consistently observed, however, that the minimum distance between the terminal domains remained above 4.0 nm, and the periodic box size was large enough to exclude any potential artifacts arising from artificial contacts or overlaps between neighboring periodic images of the domains (Figure S2).

No dissociation of EM2 from the binding pocket was observed in any set of the simulations. The orientation of EM2 stayed the same for the restrained set of simulations as expected. For the unrestrained set, except for N328^7.45^D, the ligand roughly maintained its initial position throughout the simulations, similar to that observed for the superagonist peptide DAMGO in the cryo-EM structure with regard to the spatial arrangement of pharmacophores (Figure S3). [27] The relatively large disposition of EM2 in the N328^7.45^D mutant could be due to the extra negative charge introduced in the vicinity of the ortho- and allosteric binding pockets. Interestingly, the χ^1^ torsional angle of F^3^ of EM2 adopted *gauche-* conformation dominantly, for most systems, in agreement with a previous proposal for the bioactive conformation of opioid peptides (Table S3). [66]

### 3.2. Functional dynamics

#### 3.2.1 Allosteric Na^+^ binding

The role of Na^+^ as a negative allosteric modulator for class A GPCRs, along with its allosteric binding site has been well documented, especially in ligand-free states of the receptor. [67–69] In line with the allosteric stabilization of the inactive state by Na^+^, the active-state structure of the receptors typically does not exhibit the presence of a Na^+^ ion. The allosteric Na^+^ binding site, as evidenced in previous simulations, is only accessible through the orthosteric pocket. [33–35] In the case of mutant MOP complexes, only the mutant N328^7.45^D showed Na^+^ penetration irrespective of whether the ligand was restrained or free in the binding pocket. In contrast with earlier observations, Na^+^ inserted into the N328^7.45^D mutant MOP receptor in spite of the bound ligand, indicating lower stability of the receptor ligand complex even while the ligand was mildly restrained. (Figure 2, Table 1). Intracellular access of Na^+^ ions through the TM domain is conceivably obstructed by intracellular signaling proteins, as the entrance of Na^+^ from the cytosolic side did not occur in any of the systems.

**Figure 2.**
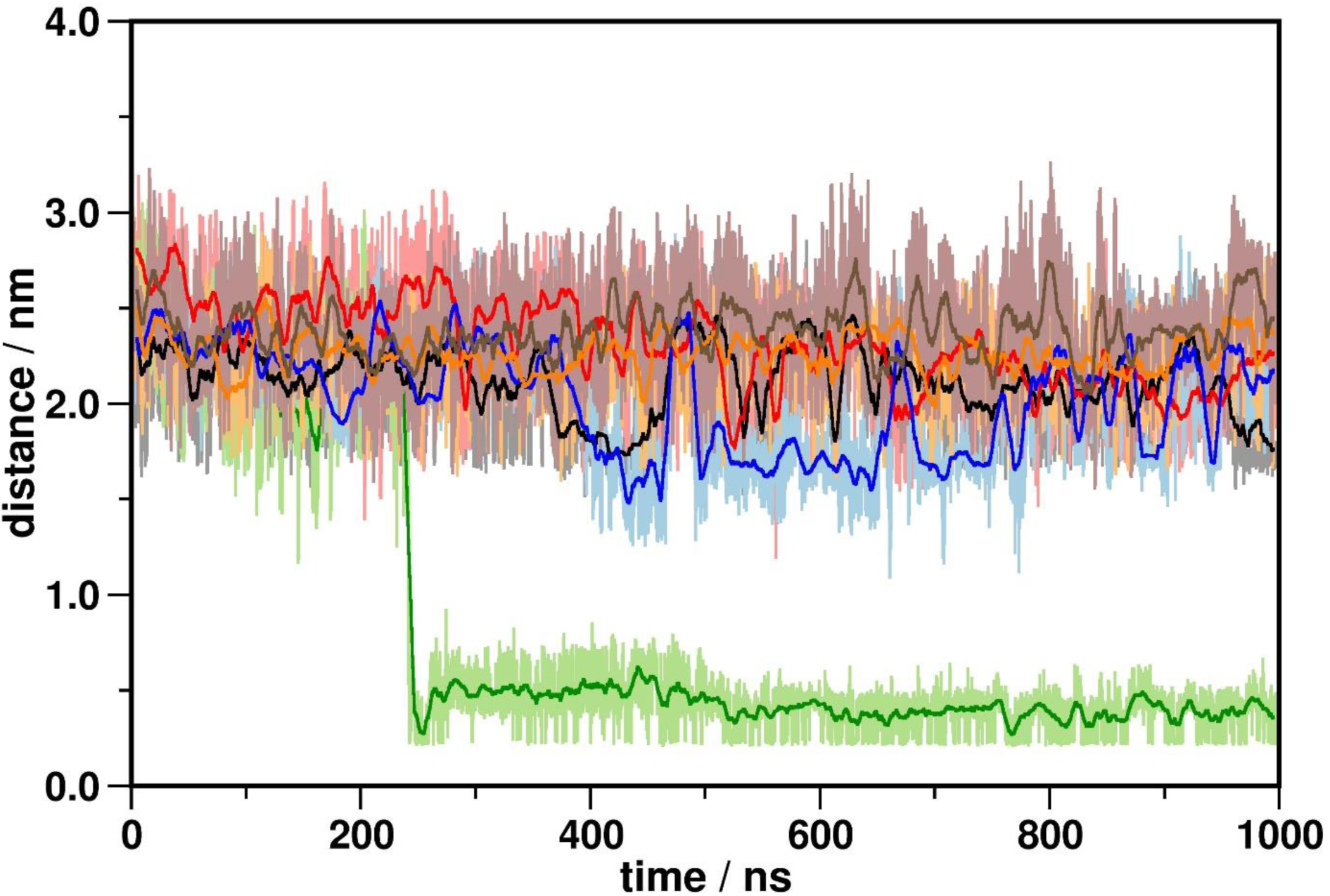
The minimum distance between Na^+^ ions and D114^2.50^ of the allosteric binding pocket Black: wild type MOP; red: Y326^7.43^F; green: N328^7.45^D; blue: N328^7.45^L; orange: D340^8.47^N; brown: D340^8.47^L.

**Table 1.**
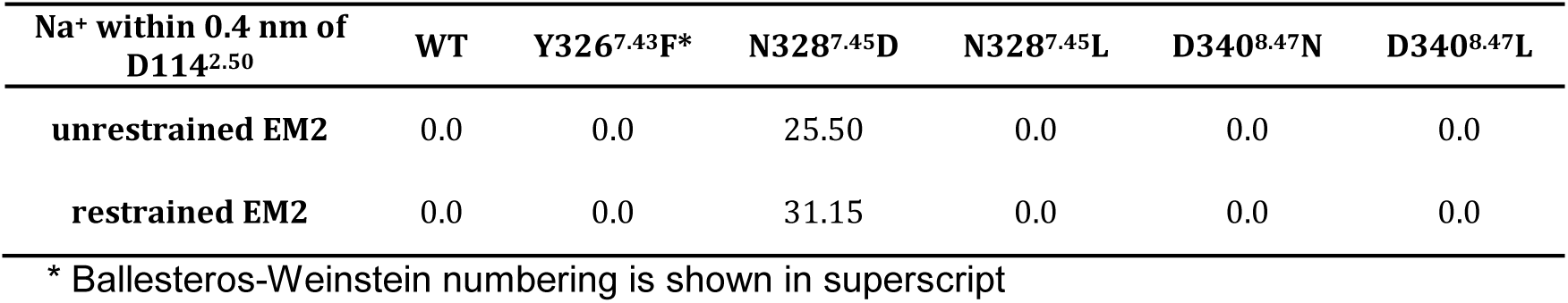
Frequency of Na^+^ ions present in the allosteric binding pocket

| Na<sup>+</sup> within 0.4 nm of D114<sup>2.50</sup> | WT | Y326<sup>7.43</sup>F* | N328<sup>7.45</sup>D | N328<sup>7.45</sup>L | D340<sup>8.47</sup>N | D340<sup>8.47</sup>L |
| --- | --- | --- | --- | --- | --- | --- |
| unrestrained EM2 | 0.0 | 0.0 | 25.50 | 0.0 | 0.0 | 0.0 |
| restrained EM2 | 0.0 | 0.0 | 31.15 | 0.0 | 0.0 | 0.0 |

\* Ballesteros-Weinstein numbering is shown in superscript

#### 3.2.2 Transmembrane helix dynamics

The disposition of TM6 in the transmembrane domain serves as a general indicator of the activation state of class A GPCRs, also referred to as the ‘signature’ conformational switch. In all simulation setups, analysis of the dynamics of TM6 helix indicated small to moderate displacements from the corresponding starting structures (Figure 3, Figure S4). TM6 maintained active-like/intermediate positions in all systems throughout the simulations and no complete transitions to inactive receptor conformations were observed. This finding aligns with previous simulation results where larger displacements of TM6 were only observed in the ligand free system, or at much longer timescales. [8,33–35] Mild restraints applied on ligand positions apparently resulted in relatively larger TM6 dispositions in some of the systems, but not to the extent of deactivation. (Figure S4)

**Figure 3.**
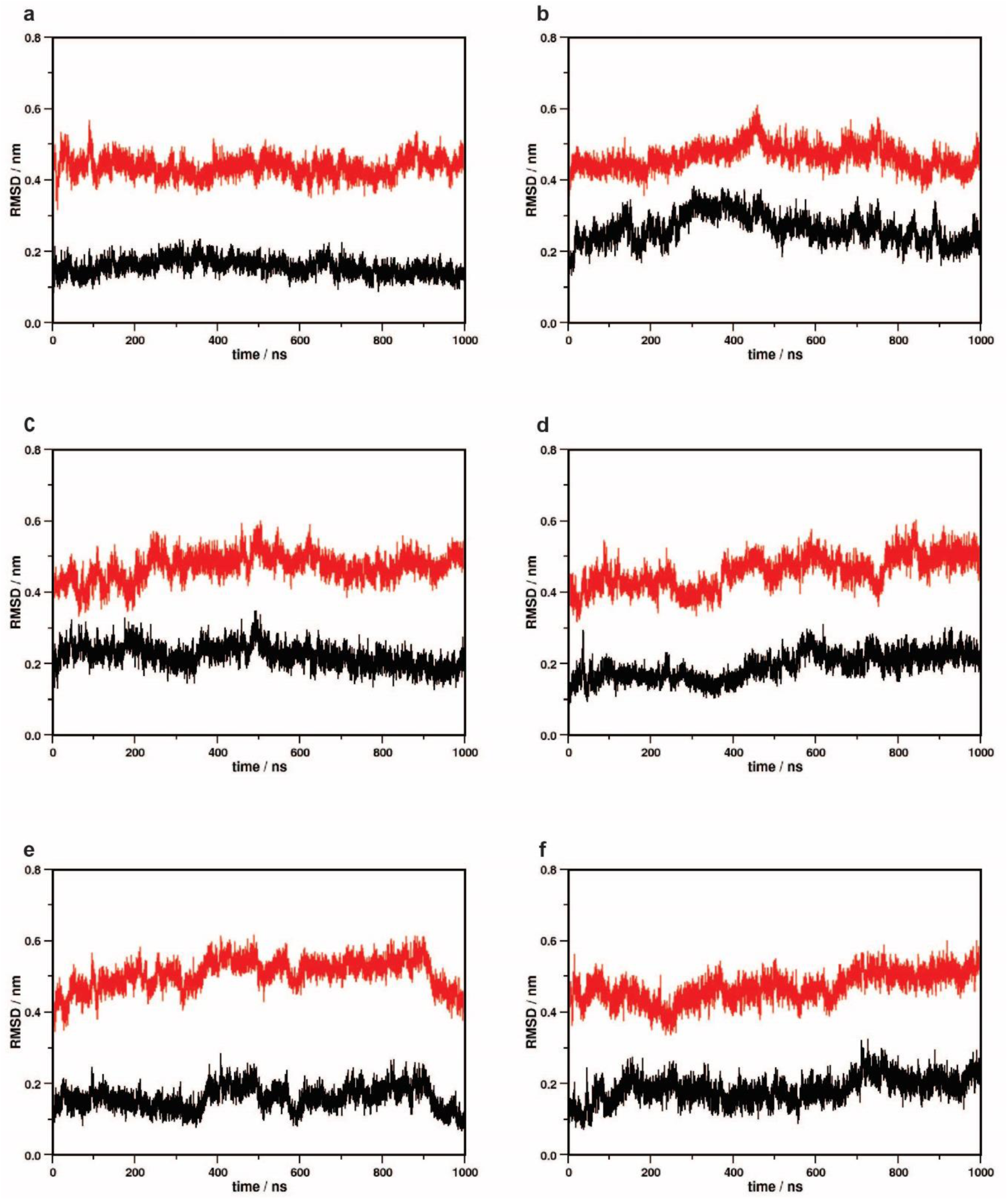
Disposition of TM6 during unrestrained simulations from the active (black) and the inactive (red) states. (a) wild type MOP (b) Y326^7.43^F (c) N328^7.45^D (d) N328^7.45^L (e) D340^8.47^N (f) D340^8.47^L.

The conserved NPxxY motif, located at the junction of TM7 and H8, is implicated in stabilizing intermediate conformations in ligand-free class A GPCRs. [8,25] These intermediate conformations are crucial for facilitating the insertion of G proteins. Notably, significant displacements of the conserved NPxxY motif were considered a contributing factor to the mobility of the polar channel segment closer to the intracellular surface, associated with the activation of the β_2_AR and type 1 cannabinoid (CB1) receptors. [34,35] The wild type MOP and its mutants generally exhibited minimal deviations of the NPxxY motif, with the exception of the N328^7.45^D mutant. The N328^7.45^D mutant displayed a substantial displacement of the motif from its initial active state position (Figure 4, Figure S5). This noteworthy observation, coupled with the reported Na^+^ penetration, suggests the profound impact that a single mutation can have on the receptor, influencing the polarity of the signaling channel preceding the NPxxY motif. The application of ligand restraints apparently promoted the mobility of this motif for most systems (Figure S5) which in the case of the N328^7.45^D mutant, similarly to the unrestrained simulations, reached the extent of structural destabilization. While in in the case of the NPxxY motif elevated mobility could be beneficial for the function of the receptor, the effect of restraints should be interpreted carefully, because locking the relative position of the ligand may also restrict the relaxation of the receptor-ligand complex, which may lead to structural destabilization of the receptor.

**Figure 4.**
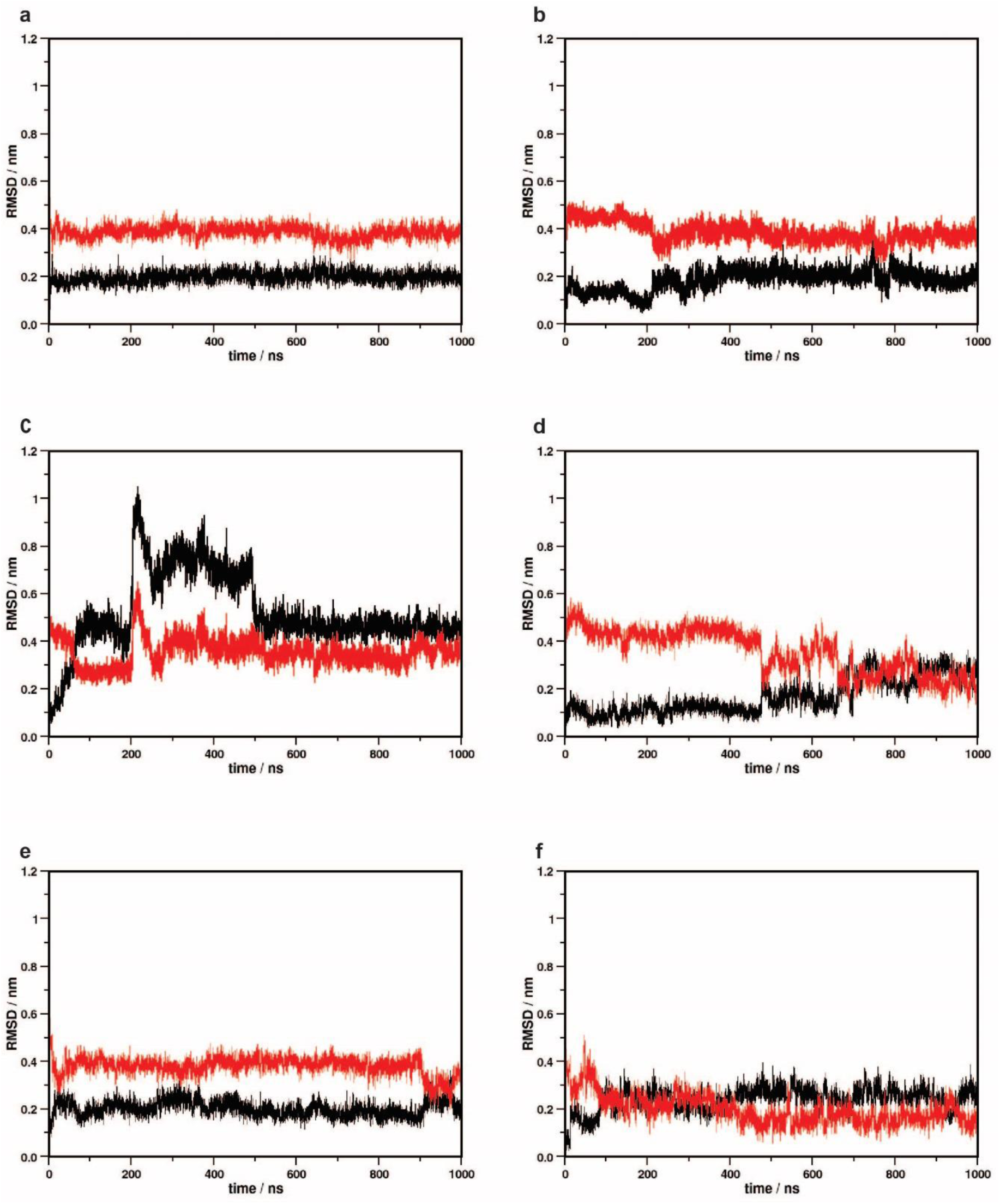
Disposition of the NPxxY motif during unrestrained simulations from the active (black) and the inactive (red) states. (a) wild type MOP (b) Y326^7.43^F (c) N328^7.45^D (d) N328^7.45^L (e) D340^8.47^N (f) D340^8.47^L.

#### 3.2.3 Intramolecular interactions

The results of MD trajectory analysis for specific intramolecular salt bridges and hydrogen bonds, previously recognized as pivotal molecular switches for class A GPCR activation, are summarized in Table 2. [25,26,36,48,70,71,72] No significant or relevant differences were found between the unrestrained and ligand restrained simulations with regard to the frequencies of these intramolecular interactions. Experimental data suggests that the salt bridge and hydrogen bond between the D^3.49^ and R^3.50^ residues of the DRY motif remain absent in most active state GPCRs. Consistent with the data, MOP and its mutants, where we dealt with just the active states, these interactions were primarily absent as expected.

**Table 2.** Frequency of specific intramolecular salt bridges and H-bonds expressed as percentages of the total conformational ensemble, generated by the unrestrained MD simulations.

| Interactions | Residues involved* | WT | Y326<sup>7.43</sup>F | N328<sup>7.45</sup>D | N328<sup>7.45</sup>L | D340<sup>8.47</sup>N | D340<sup>8.47</sup>L |
| --- | --- | --- | --- | --- | --- | --- | --- |
| **Salt bridges** |  |  |  |  |  |  |  |
| DRY – H8 | R165<sup>3.50</sup>, D340<sup>8.47</sup> | 7.45 | 23.66 | 1.26 | 0.53 | N/A | N/A |
| intra-DRY | D164<sup>3.49</sup>, R165<sup>3.50</sup> | 0.00 | 0.00 | 0.00 | 0.02 | 0.02 | 0.00 |
| **H-bonds** |  |  |  |  |  |  |  |
| DRY - TM6 | R165<sup>3.50</sup>, T279<sup>6.34</sup> | 0.00 | 0.00 | 0.00 | 0.00 | 0.00 | 0.00 |
| DRY - ICL2 | D164<sup>3.49</sup>, R179<sup>ICL2</sup> | 99.52 | 100.00 | 100.00 | 99.99 | 92.36 | 99.94 |
| DRY - TM5 | R165<sup>3.50</sup>, Y252<sup>5.58</sup> | 9.08 | 0.00 | 80.96 | 5.11 | 40.07 | 9.72 |
| CWxP - TM7 | C292<sup>6.47</sup>, W293<sup>6.48</sup>, N328<sup>7.45</sup> | 0.08 | 0.12 | 0.01 | 0.00 | 27.37 | 0.03 |
| NPxxY- TM network | N332<sup>7.49</sup>, Y336<sup>7.53</sup>, L158<sup>3.43</sup>, Y252<sup>5.58</sup>, V285<sup>6.40</sup> | 9.86 | 42.09 | 2.93 | 16.21 | 1.53 | 0.00 |

\* Ballesteros-Weinstein numbering is shown in superscript

Our simulation results supported the concept of an ’ionic lock’ interaction between the DRY motif and TM6. This interaction was proposed to serve as a constraint in the inactive state, and its disruption upon receptor activation was suggested to facilitate the outward movement of TM6. This is an exclusive property for the inactive state since none of the active state MOP and its mutants showed the presence of the ‘ionic lock’ during the simulations. The identification of a distinctive salt bridge between R165^3.50^ of the DRY motif and D340^8.47^ of H8 was initially highlighted in our previous MD simulation study of the MOP. [33] However, in our current study, as well as in other previous studies of the β_2_AR and the CB1 receptors such interactions mostly remained absent, and could be deemed as transient, observed only in that particular case.

H-bonds were systematically present between D164^3.49^ of the DRY motif and R179 of the second intracellular loop (ICL2) in all MOP variants. This interaction was suggested previously to reorganize upon the activation of the β_2_AR [73], but that was not confirmed by our previous MD simulation studies of that particular receptor. [34] Here, minor structural destabilization of ICL2 of the Y326^7.43^F was found (Figure S6), but no clear correlation of this exception with the other analysis results is present that could justify the relevance of this observation. Furthermore, no clear trends could be identified for the frequencies of the CWxP-TM7 and NPxxY-TM network interactions. The DRY-TM5 interaction, proposed earlier to stabilize the G protein-bound active state was most frequently found in the N328^7.45^D mutant, which is in clear contradiction with the above presented results and inferences regarding the stability of this mutant receptor ligand complex and the integrity of its signaling state structure. These interaction patterns, observed originally in specific experimental GPCR structures, are not generally applicable as they were shown previously to differ largely in different GPCRs. [33–35]

#### 3.2.4 Correlated dynamics among polar signaling channel residues

The observation of the correlated motions among the polar signaling channel residues, indicated by GCCM analysis (Figure 5, Figure S7-S8), has been consistent in the active, G protein bound state of the receptors, in our earlier as well as current studies. However, there are few differences among their dynamics which could be attributed to the specific properties of the mutant receptors. Figures 6 and S9 show the schematic representation of correlated motions observed between the channel residues of MOP and its mutants during the simulations. The complete correlation of movements of the polar signaling channel residues, pertaining to receptor activation and consequent signaling were reproduced in the wild type receptor system in both setups, as well as in the D340^8.47^N mutant simulation, when the ligand was not restrained. Other mutant systems failed to show such absolute synchronous movement among the polar channel residues, suggesting that the only mutation that is possibly tolerated is the one which is distant from the orthosteric and allosteric binding pockets and in which the polarity is preserved. The involvement of D114^2.50^ in the correlated motions was particularly missing in these observations, underscoring the impact of D114^2.50^**-**N332^7.49^ internal dynamics in the activation and signaling of MOP. [33,40] The ordered dynamics of the polar signaling channel, observed in the wild type and the D340^8.47^N mutant MOP receptors, is presumed to propagate the activation signal in the form of charge perturbation towards the G protein coupling interface, assisted by water molecules present in the cavity. The presence of a hydrated pathway along the polar channel, with rapid exchange among the water molecules has been consistently observed in all relevant systems (Figure S10), and plays a role in signal transmission in the active receptor complexes. [14,74]

**Figure 5.**
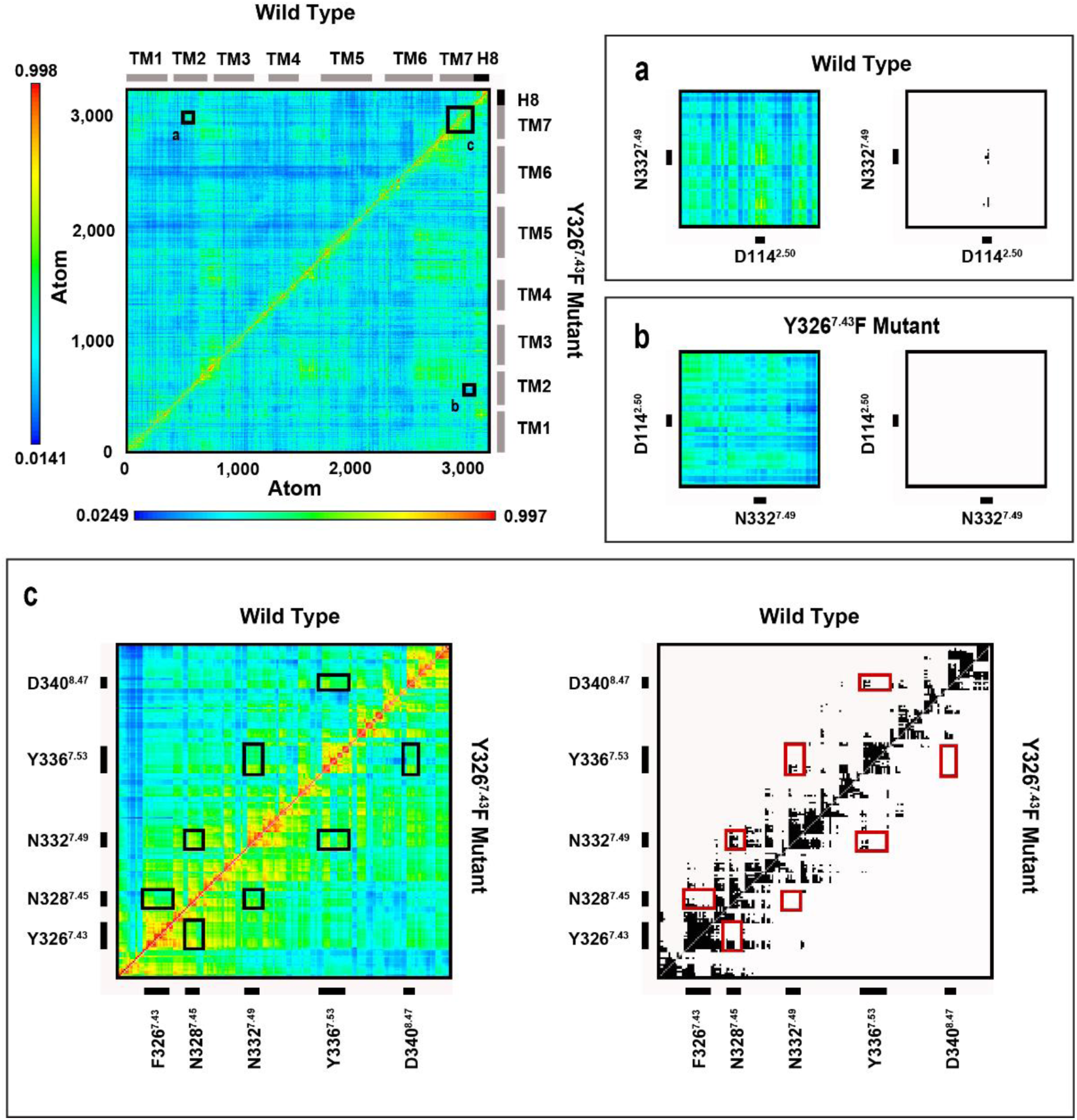
Dynamic cross-correlation matrices of the active state, G_i_ protein-bound D340^8.47^N and D340^8.47^L MOP mutants. Panels (a-c) are magnified views of regions of amino acid residues of interest. Black and white panels show correlations above the threshold of 0.63 MI.

**Figure 6.**
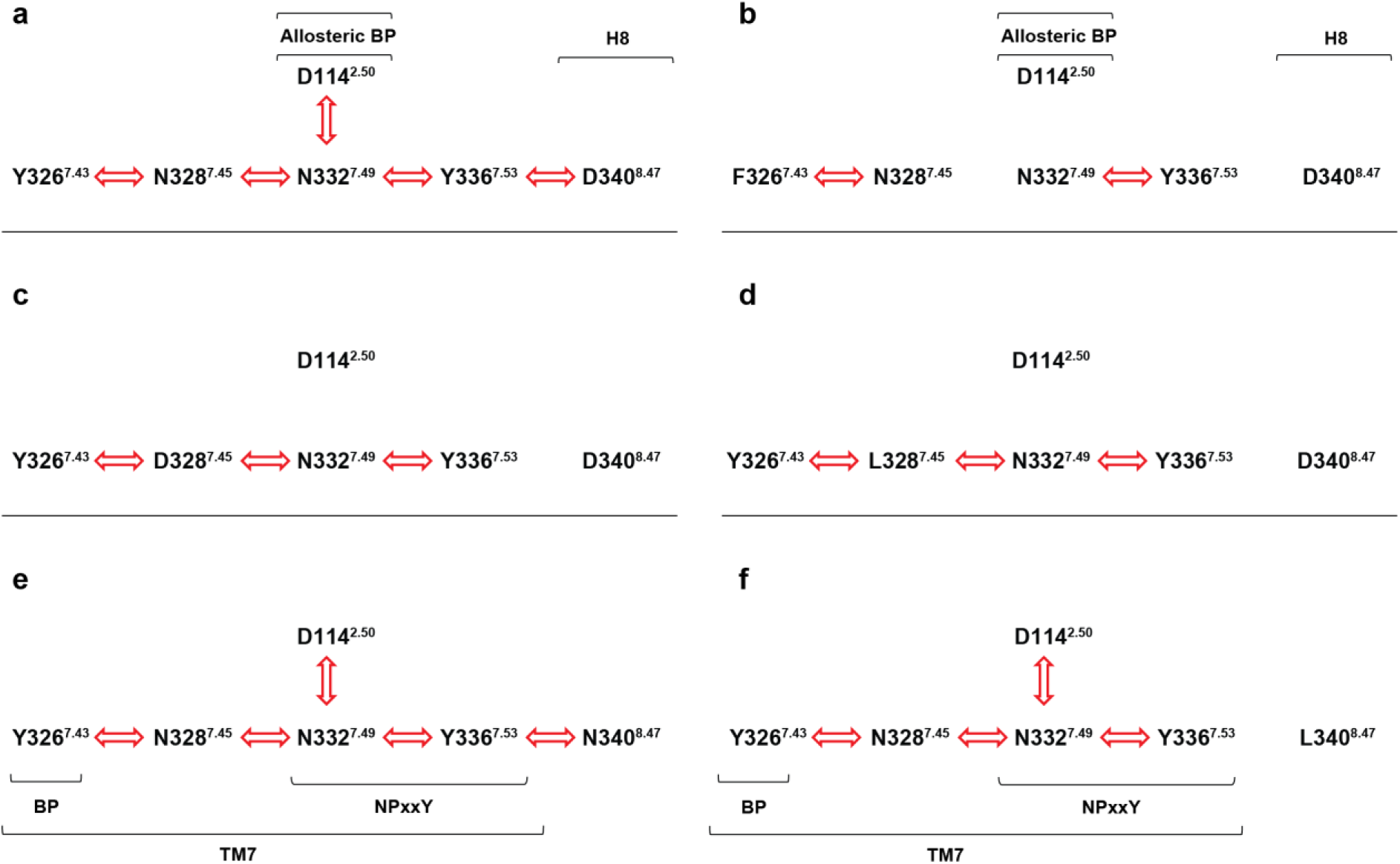
Correlated motions in the conserved polar signaling channel of active state, G_i_ protein-bound MOP derivatives revealed by generalized cross correlation matrix analysis (unrestrained simulations). Red arrows indicate correlated motions of the respective amino acids in the (a) wild type MOP (b) Y326^7.43^F (c) N328^7.45^D (d) N328^7.45^L (e) D340^8.47^N (f) D340^8.47^L mutants.

The results provide support to our hypothesis, that altering the polarity of the constituents of the signaling channel does affect the dynamics between the polar side chains, subsequently has a detrimental effect on signal transduction. Another important takeaway could be extended to the orientation of the ligand during the course of the simulation. The primary requisite is the starting orientation of the ligand to be in its bioactive conformation. However, forced unnatural restraint applied to fix EM2 even in its bioactive orientation, could result in the restriction of ordered movement. It can be presumed that ligand movement within the orthosteric pocket has to be well coordinated, and even a minor deviation from it could lead to the arrest of signal propagation initiated from this site.

#### 3.2.5 Pharmacological assessment

The results of *in vitro* radioligand competition binding experiments demonstrated, that EM2 was able to fully displace the radiolabeled prototypic agonist [^3^H]DAMGO from the orthosteric pocket the wild type MOP and the D340^8.47^N mutant. This was further corroborated by the comparable binding affinities, with an inhibition constant of 2.0 nM and 1.4 nM for the wild type and the mutant MOP, respectively, in good agreement with previously published data [64]. Conversely, no orthosteric interaction between EM2 and the rest of the mutant receptors was observed (Figure 7a, Table 3)

**Figure 7.**
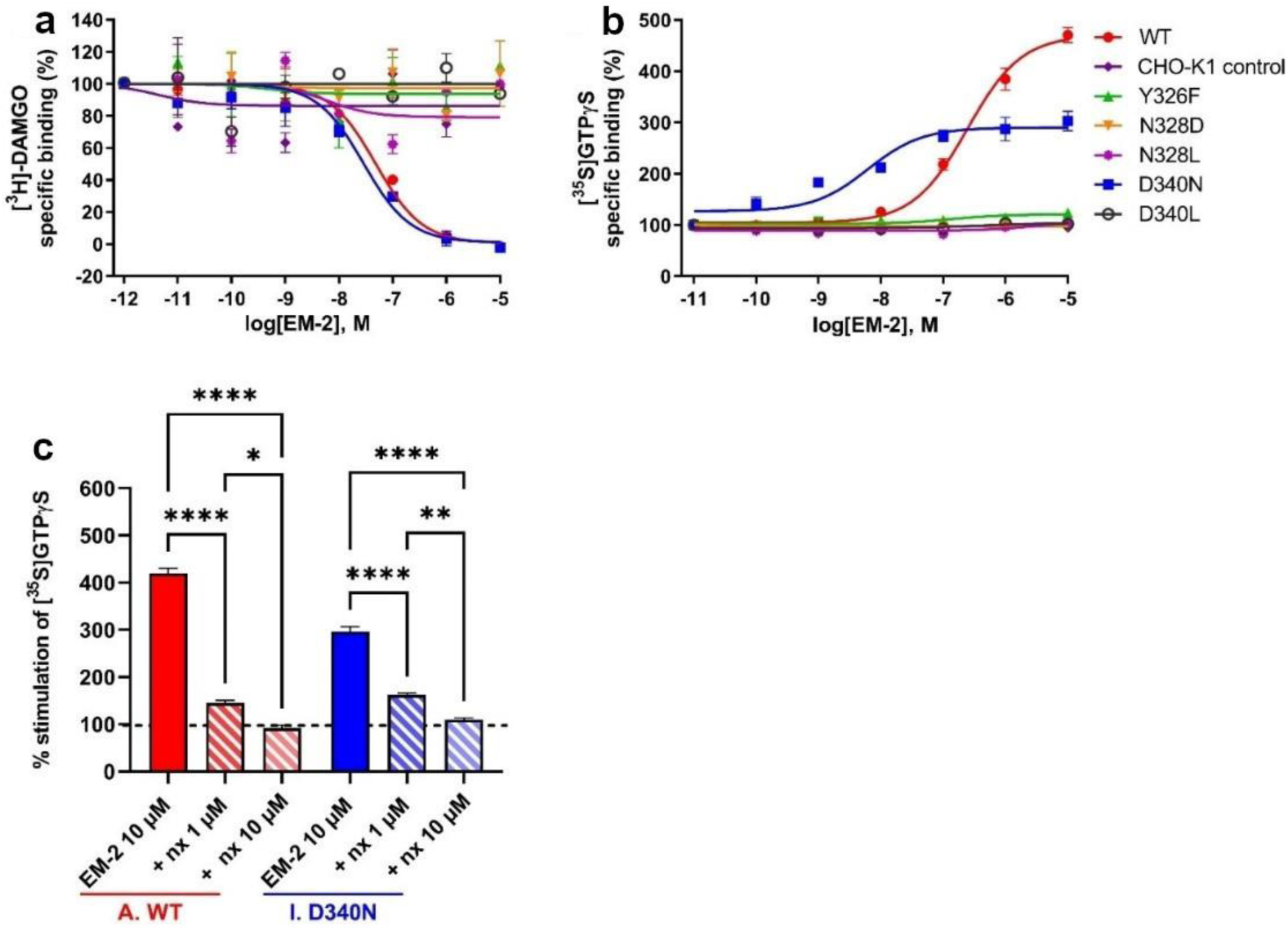
(a) MOP receptor binding affinity of EM2 in [^3^H]DAMGO competition binding assays to cell membrane homogenates. Figures represent the specific binding of the radioligand in percentage in the presence of increasing concentrations (10^−12^–10^−5^ M) of the indicated ligands. Data are expressed as percentage of mean specific binding ± S.E.M. (n ≥ 3). (b) G protein activation effects of Endomorphin-2 ligand in [^35^S]GTPγS binding assays in cell membrane homogenates. Figures represent the relative specific binding of [^35^S]GTPγS in the presence of increasing concentrations (10^−11^–10^−5^ M) of the indicated compound, EM2. Data are expressed as percentage of mean specific binding ±S.E.M. (n ≥ 3) over the basal activity (100%), (C) dose-dependent (1 and 10 µM) inhibition of 10 µM endomorphin-2 (EM-2) [^35^S]GTPγS binding by the MOP antagonist, naloxone (nx) in wild type and D340^8.47^N mutant cell membrane homogenates. 100% is the basal activity. Data are expressed as a percentage of mean specific binding ± S.E.M. (n ≥ 3). ****, P < 0.0001, **, P < 0.01, *, P < 0.05, one-way ANOVA followed by Tukey’s multiple comparison test.

**Table 3.** Binding affinity (K_i_) and signaling efficacy (E_max_) and potency (EC_50_) of the MOP selective endogenous peptide EM2 on membranes of CHO-K1 cells stably expressing the wild type and mutant MOP.

| Cells | EM2 vs [$^3$H]-DAMGO | EM2 stimulated<br>[$^{35}$S]GTPγS binding | |
| --- | --- | --- | --- |
| | $K_i \pm$ S.E.M. (nM) | $E_{\max} \pm$ S.E.M. (%) | $EC_{50} \pm$ S.E.M. (nM) |
| CHO-K1 MOP WT | $2 \pm 0.3$ | $437 \pm 9.7^{****}$ | $245 \pm 16^{***}$ |
| CHO-K1 negative control | n.b. | $94 \pm 4.1$ | n.r |
| CHO-K1 MOP Y326<sup>7.43</sup>F | n.b. | $113 \pm 6.6$ | n.r |
| CHO-K1 MOP N328<sup>7.45</sup>D | n.b. | $101 \pm 2.8$ | n.r |
| CHO-K1 MOP N328<sup>7.45</sup>L | n.b. | $107 \pm 2.6$ | n.r |
| CHO-K1 MOP D340<sup>8.47</sup>N | $1.4 \pm 0.07$ | $290 \pm 9.8^{****}$ | $11 \pm 4^{***}$ |
| CHO-K1 MOP D340<sup>8.47</sup>L | n.b. | $104 \pm 2.8$ | n.r |

EM-2: MOP agonist; n.b.: no binding. n.r.: not relevant $n \geq 3$ . Statistical comparison of stimulation efficacy ( $E_{\max}$ ) was performed by one-way ANOVA followed by Tukey's multiple comparison test (\*\*\*\*, $P < 0.0001$ ). \*\*\*, significant difference ( $P < 0.05$ ) from ligand potency ( $EC_{50}$ ) values were determined by unpaired Student's t-test, in the binding affinity ( $K_i$ ) values there was no significance.

When signaling potency of the wild type and mutant receptors was examined through agonist-stimulated [^35^S]GTPγS binding functional assays, it was observed that in both the wild type and the D340^8.47^N mutant MOP EM2 induced significant alterations in basal [^35^S]GTPγS binding which indicated G protein activation. (Figure 7b, Table 3) Although EM2 exhibited a full agonistic effect on both receptors, there were notable differences in efficacy (E_max_) and potency (EC_50_) values. For the wild-type MOP, EM2 efficiently stimulated G proteins with high efficacy (437%) and moderate potency (245 nM). In the D340^8.47^N mutant, EM2 demonstrated lower efficacy (290%) but increased potency (11 nM). These variations in the ligand’s stimulatory effect suggest that the D340^8.47^N mutation does not diminish, yet affects receptor functionality. Consistent with the results of EM2 binding experiments, the other mutants failed to exhibit any functional activity. The agonist effect of EM2 was demonstrated to be dose-dependently inhibited by the clinically used MOP antagonist, naloxone, in both wild type and D340^8.47^N mutant receptors. The results indicated that nearly all mutant systems led to the complete loss of both ligand binding and signaling capability. D340^8.47^N as mentioned, preserved or maintained the polarity in the channel. This observation perfectly corroborated our findings from simulation studies, where this remained the only mutant showing complete correlated motions among the polar side chains. Another particularly significant observation was that a radical mutation in the intracellular G protein coupling region (D340^8.47^L) resulted in impaired ligand binding at the orthosteric region (Figure 1, Figure 7, Table 3). This outcome provides direct support to the hypothesis of a strong allosteric linkage between these two sites crucial for receptor activation.

## 4. Conclusions

The existence of interconnecting microswitches extending from the ligand binding pocket to the intracellular surface, and its prime importance in the activation and signaling of class A GPCRs has been well established already in experiments as well as computational studies. [15,20] However, most of the discourse was based on large-scale structural rearrangements of the TM domain and associated macroscopic changes. These also give in to the bottleneck formed in situations, when multiple active states of the receptor or structurally analogous but functionally different ligands are considered and no sufficiently accurate explanation for the activation of different signaling pathways could be given based on structural rearrangements alone. As proposed earlier, an additional event underlying these phenomena could be a shift in the charge balance and the consequent polarization of the receptor. Such polarization could be contributed by the concerted motions of polar species in the TM domain, as well as the nature of the orthosteric ligand. [33–35]. GCCM analysis confirmed such concerted motions in the polar signaling channel in the wild type, as well as the D340^8.47^N mutant receptors, but not in the other mutants, indicating that significant alteration of the polarity of channel constituents is detrimental for both the ligand binding and the signaling capacity of the MOP. The bioactive orientation of the ligand and its native free dynamics in the orthosteric pocket was also suggested to be essential for receptor activation. The conclusions drawn from the simulation data were corroborated by results of *in vitro* competitive radioligand binding assays as well as [^35^S]GTPγS binding functional assays. Furthermore, experimental results provided direct evidence for strong allosteric coupling between the orthosteric binding pocket and the intracellular surface of the MOP. The results obtained from our current study along with earlier findings provide further insights into the activation pathways of class A GPCRs which could have important implications for the development of new therapeutic strategies targeting this receptor family. More specifically, the new perspective introduced, concerning the electrostatic balance of the TM domain could open new avenues in the development of novel, pathway specific drugs.

## Author Contributions

Conceptualization, A.B.; methodology, A.S., D.Sz., A.M., L.Z. H.M., K.B. and A.B.; formal analysis, A.S. and D.Sz.; investigation, A.S. and D.Sz.; writing - original draft preparation, A.S. and A.B.; writing - review and editing, A.S., A.B., Z.L. H.M., D.Sz and K.B.; visualization, A.S. and A.B.; supervision, A.B.; project administration, A.B. All authors have read and agreed to the published version of the manuscript.

## Institutional Review Board Statement

Not applicable.

## Informed Consent Statement

Not applicable.

## Data Availability Statement

All data presented in the article and the Supplementary Materials are available upon request from the corresponding author.

## Acknowledgments

Computing resources were provided by the Government Agency of Information Technology Development, Hungary. The authors are grateful to Katalin Udvardy and Géza Tóth (Institute of Biochemistry, HUN-REN Biological Research Centre, Szeged, Hungary) for assistance with the cell cultures and for providing synthetic EM2, respectively. The authors also would like to extend their gratitude to András Váradi (Memorial Sloan-Kettering Cancer Center, New York, United States) for supplying the CHO-MOP cell lines.

## Abbreviations

cryo-EM: cryo-electron microscopy
EM: endomorphin
GCCM: generalized cross correlation matrix
GDP: guanosine diphosphate
GPCR: G protein-coupled receptor
GTP: guanosine triphosphate
ICL: intracellular loop
MD: molecular dynamics
MOP: μ-opioid receptor
PTM: post-translational modification
TM: transmembrane

## References

1. Hauser AS, Attwood MM, Rask-Andersen M, Schiöth HB, Gloriam DE. Trends in GPCR drug discovery: new agents, targets and indications. Nature Reviews Drug Discovery 16, 829–842 (2017). DOI:10.1038/nrd.2017.178

2. Santos R, Ursu O, Gaulton A, Bento AP, Donadi RS, et al. A comprehensive map of molecular drug targets. Nature Reviews Drug Discovery 16, 19–34 (2016). DOI:10.1038/nrd.2016.230

3. Childers SR. Opioid receptor-coupled second messenger systems. Life Sciences 48, 1991–2003 (1991). DOI:10.1016/0024-3205(91)90154-4

4. Schmidt H, Schulz S, Klutzny M, Koch T, Händel M, Höllt V. Involvement of Mitogen-Activated Protein Kinase in Agonist-Induced Phosphorylation of the μ-Opioid Receptor in HEK 293 Cells. Journal of Neurochemistry 74, 414–422 (2000). DOI:10.1046/j.1471-4159.2000.0740414.x

5. Xie W, Samoriski GM, McLaughlin JP, Romoser VA, Smrcka A, et al. Genetic alteration of phospholipase C β3 expression modulates behavioral and cellular responses to μ opioids. Proceedings of the National Academy of Sciences 96, 10385–10390 (1999). DOI:10.1073/pnas.96.18.10385

6. Pándy-Szekeres G, Munk C, Tsonkov TM, Mordalski S, Harpsøe K, Hauser AS, Bojarski AJ, Gloriam DE. GPCRdb in 2018: adding GPCR structure models and ligands. Nucleic Acids Research 46, 440-446 (2017). DOI:10.1093/nar/gkx1109

7. Latorraca NR, Venkatakrishnan AJ, Dror RO. GPCR Dynamics: Structures in Motion. Chemical Reviews 117, 139–155 (2017). DOI:10.1021/acs.chemrev.6b00177

8. Dror RO, Arlow DH, Maragakis P, Mildorf TJ, Pan AC, et al. Activation mechanism of the *β*_2_-adrenergic receptor. Proceedings of the National Academy of Sciences 108, 18684–18689 (2011). DOI:10.1073/pnas.1110499108

9. Kapoor A, Martinez-Rosell G, Provasi D, de Fabritiis G, Filizola, M. Dynamic and Kinetic Elements of µ-Opioid Receptor Functional Selectivity. Scientific Reports 7, 11255 (2017). DOI:10.1038/s41598-017-11483-8

10. Manglik A, Kruse AC. Structural Basis for G Protein-Coupled Receptor Activation. Biochemistry 56, 5628–5634 (2017). DOI:10.1021/acs.biochem.7b00747

11. Marino KA, Prada-Gracia D, Provasi D, Filizola M. Impact of Lipid Composition and Receptor Conformation on the Spatio-temporal Organization of μ-Opioid Receptors in a Multi-component Plasma Membrane Model. PLoS Computational Biology 12, e1005240 (2016). DOI:10.1371/journal.pcbi.1005240

12. Strohman MJ, Maeda S, Hilger D, Masureel M, Du Y, Kobilka BK. Local membrane charge regulates β_2_ adrenergic receptor coupling to G_i3_. Nature Communications 10, 2234 (2019). DOI:10.1038/s41467-019-10108-0

13. Pert CB, Pasternak G, Snyder SH. Opiate agonists and antagonists discriminated by receptor binding in brain. Science 182, 1359–1361 (1973). DOI:10.1126/science.182.4119.1359

14. Venkatakrishnan AJ, Ma AK, Fonseca R, Latorraca NR, Kelly B, et al. Diverse GPCRs exhibit conserved water networks for stabilization and activation. Proceedings of the National Academy of Sciences 116, 3288–3293 (2019). DOI:10.1073/pnas.1809251116

15. Filipek S. Molecular switches in GPCRs. Current Opinion in Structural Biology 55, 114–120 (2019). DOI:10.1016/j.sbi.2019.03.017

16. Liu W, Chun E, Thompson AA, Chubukov P, Xu F, et al. Structural Basis for Allosteric Regulation of GPCRs by Sodium Ions. Science 337, 232–236 (2012). DOI:10.1126/science.1219218

17. Fenalti G, Giguere PM, Katritch V, Huang XP, Thompson AA, et al. Molecular control of δ-opioid receptor signalling. Nature 506, 191–196 (2014). DOI:10.1038/nature12944

18. Marino KA, Shang Y, Filizola M. Insights into the function of opioid receptors from molecular dynamics simulations of available crystal structures. British Journal of Pharmacology 175, 2834–2845 (2018). DOI:10.1111/bph.13774

19. Katritch V, Fenalti G, Abola EE, Roth BL, Cherezov V, Stevens RC. Allosteric sodium in class A GPCR signaling. Trends in Biochemical Sciences 39, 233–244 (2014). DOI:10.1016/j.tibs.2014.03.002

20. Zhou Q, Yang D, Wu M, Guo Y, Zhong L, et al. Common activation mechanism of class A GPCRs. eLife 8, e50279 (2019). DOI:10.7554/eLife.50279

21. Qu Q, Huang W, Aydin D, Paggi JM, Seven AB, et al. Insights into distinct signaling profiles of the µOR activated by diverse agonists. Nature Chemical Biology 19, 423–430 (2023). DOI:10.1038/s41589-022-01208-y

22. Suomivuori CM, Latorraca NR, Wingler LM, Eismann S, King MC, et al. Molecular mechanism of biased signaling in a prototypical G protein–coupled receptor. Science 367, 881–887 (2020). DOI:10.1126/science.aaz0326

23. Williams JT, Ingram ST, Henderson G, Chavkin C, von Zastrow M, et al. Regulation of *μ*-Opioid Receptors. Pharmacological Research 65, 223–254 (2013). DOI:10.1124/pr.112.005942

24. Waldhoer M, Bartlett SE, Whistler JL. Opioid Receptors. Annual Review of Biochemistry 73, 953–990 (2004). DOI:10.1146/annurev.biochem.73.011303.073940

25. Huang W, Manglik A, Venkatakrishnan AJ, Laeremans T, Feinberg EN, et al. Structural insights into µ-opioid receptor activation. Nature 524, 315–321 (2015). DOI:10.1038/nature14886

26. Manglik A, Kruse AC, Kobilka TS, Thian FS, Mathiesen JM, et al. Crystal structure of the μ-opioid receptor bound to a morphinan antagonist. Nature 485, 321–326 (2012). DOI:10.1038/nature10954

27. Koehl A, Hu H, Maeda S, Zhang Y, Qu Q, et al. Structure of the μ Opioid Receptor-Gi Protein Complex. Nature 558, 547–552 (2018). DOI:10.1038/s41586-018-0219-7

28. Wang Y, Zhuang Y, DiBerto JF, Zhou XE, Schmitz GP, et al. Structures of the entire human opioid receptor family. Cell 186, 413–427 (2023). DOI:10.1016/j.cell.2022.12.026

29. Faouzi A, Wang H, Zaidi SA, DiBerto JF, Che T, et al. Structure-based design of bitopic ligands for the µ-opioid receptor. Nature 613, 767–774 (2023). DOI:10.1038/s41586-022-05588-y

30. Zhuang Y, Wang Y, He B, He X, Zhou X.E, et al. Molecular recognition of morphine and fentanyl by the human μ-opioid receptor, Cell 185, 4361–4375.e19 (2022). DOI:10.1016/j.cell.2022.09.041

31. Wang H, Hetzer F, Huang W, Qu Q, Meyerowitz J, et al. Structure-Based Evolution of G Protein-Biased μ-Opioid Receptor Agonists. Angewandte Chemie International Edition 61, e202200269 (2022). DOI:10.1002/anie.202200269

32. Robertson MJ, Papasergi-Scott MM, He F, Seven AB, Meyerowitz JG, et al. Structure determination of inactive-state GPCRs with a universal nanobody. Nature Structural & Molecular Biology 29, 1188–1195 (2022). DOI:10.1038/s41594-022-00859-8

33. Mitra A, Sarkar A, Szabó MR, Borics A. Correlated Motions of Conserved Polar Motifs Lay out a Plausible Mechanism of G Protein-Coupled Receptor Activation. Biomolecules 11, 670 (2021). DOI:10.3390/biom11050670

34. Mitra A, Sarkar A, Borics A. Universal Properties and Specificities of the β_2_-Adrenergic Receptor-G_s_ Protein Complex Activation Mechanism Revealed by All-Atom Molecular Dynamics Simulations. International Journal of Molecular Sciences 22, 10423 (2021). DOI:10.3390/ijms221910423

35. Sarkar A, Mitra A, Borics A. All-Atom Molecular Dynamics Simulations Indicated the Involvement of a Conserved Polar Signaling Channel in the Activation Mechanism of the Type I Cannabinoid Receptor. International Journal of Molecular Sciences 24, 4232 (2023). DOI:10.3390/ijms24044232

36. Jongejan A, Bruysters M, Ballesteros JA, Haaksma E, Bakker RA, et al. Linking Agonist Binding to Histamine H1 Receptor Activation. Nature Chemical Biology 1, 98–103 (2005). DOI:10.1038/nchembio714

37. Liu R, Nahon D, le Roy B, Lenselink EB, Ijzerman AP. Scanning Mutagenesis in a Yeast System Delineates the Role of the NPxxY(x)5,6F Motif and Helix 8 of the Adenosine A2B Receptor in G Protein Coupling. Biochemical Pharmacology 95, 290–300 (2015). DOI:10.1016/j.bcp.2015.04.005

38. Hothersall JD, Torella R, Humphreys S, Hooley M, Brown A, et al. Residues W320 and Y328 within the Binding Site of the μ-Opioid Receptor Influence Opiate Ligand Bias. Neuropharmacology 118, 46–58 (2017). DOI:10.1016/j.neuropharm.2017.03.007

39. Sealfon SC, Chi L, Ebersole BJ, Rodic V, Zhang D, et al. Related Contribution of Specific Helix 2 and 7 Residues to Conformational Activation of the Serotonin 5-HT2A Receptor. Journal of Biological Chemistry 270, 16683–16688 (1995). DOI:10.1074/jbc.270.28.16683

40. Xu W, Ozdener F, Li JG, Chen C, de Riel JK, et al. Functional Role of the Spatial Proximity of Asp114^2.50^ in TMH 2 and Asn332^7.49^ in TMH 7 of the Mu Opioid Receptor. FEBS Letters 447, 318–324(1999). DOI:10.1016/S0014-5793(99)00316-6

41. Galés C, Kowalski-Chauvel A, Dufour MN, Seva C, Moroder L, et al. Mutation of Asn-391 within the Conserved NPxxY Motif of the Cholecystokinin B Receptor Abolishes Gq Protein Activation without Affecting Its Association with the Receptor. Journal of Biological Chemistry 275, 17321–17327 (2000). DOI:10.1074/jbc.M909801199

42. Kalatskaya I, Schüssler S, Blaukat A, Müller-Esterl W, Jochum M, et al. Mutation of Tyrosine in the Conserved NPxxY Sequence Leads to Constitutive Phosphorylation and Internalization, but Not Signaling of the Human B2 Bradykinin Receptor. Journal of Biological Chemistry 279, 31268–31276 (2004). DOI:10.1074/jbc.M401796200

43. Prioleau C, Visiers I, Ebersole BJ, Weinstein H, Sealfon SC. Conserved Helix 7 Tyrosine Acts as a Multistate Conformational Switch in the 5-HT2C Receptor. Identification of a Novel “LOCKED-ON” Phenotype and Double Revertant Mutations. Journal of Biological Chemistry 277, 36577–36584 (2002). DOI:10.1074/jbc.M206223200

44. Hunyadi L, Bor M, Baukal AJ, Balla T, Catt KJ. A Conserved NPLFY Sequence Contributes to Agonist Binding and Signal Transduction but Is Not an Internalization Signal for the Type 1 Angiotensin II Receptor. Journal of Biological Chemistry 270, 16602–16609 (1995). DOI:10.1074/jbc.270.28.16602

45. Mansour A, Taylor LP, Fine JL, Thompson RC, Hoversten MT, et al. Key Residues Defining the μ-Opioid Receptor Binding Pocket: A Site-Directed Mutagenesis Study. Journal of Neurochemistry 68, 344–353 (2002). DOI:10.1046/j.1471-4159.1997.68010344.x

46. Wall MA, Coleman DE, Lee E, Iñiguez-Lluhi JA, Posner BA, et al. The structure of the G protein heterotrimer G_iα1_β_1_γ_2_. Cell 83, 1047–1058 (1995). DOI:10.1016/0092-8674(95)90220-1

47. Jo S, Kim T, Iyer VG, Im W. CHARMM-GUI: A Web-based Graphical User Interface for CHARMM. Journal of Computational Chemistry 29, 1859–1865 (2008). DOI:10.1002/jcc.20945

48. Rasmussen SGF, DeVree BT, Zou Y, Kruse AC, Chung KY, et al. Crystal structure of the β_2_ adrenergic receptor–Gs protein complex. Nature 477, 549–555 (2011). DOI:10.1038/nature10361

49. Abraham MJ, Murtola T, Schulz R, Páll S, Smith JC, et al. GROMACS: High performance molecular simulations through multi-level parallelism from laptops to supercomputers. SoftwareX 1-2, 19-25 (2015). DOI:10.1016/j.softx.2015.06.001

50. Li DW, Brüschweiler R. NMR-Based Protein Potentials. Angewandte Chemie 49, 6778–6780 (2010). DOI:10.1002/anie.201001898

51. Qiu D, Shenkin PS, Hollinger FP, Still WC. The GB/SA Continuum Model for Solvation. A Fast Analytical Method for the Calculation of Approximate Born Radii. The Journal of Physical Chemistry A 101, 3005–3014 (1997). DOI:10.1021/jp961992r

52. Johansson MU, Zoete V, Michielin O, Guex N. Defining and searching for structural motifs using DeepView/Swiss-PdbViewer. BMC Bioinformatics 13, (2012). DOI:10.1186/1471-2105-13-173

53. Bussi G, Donadio D, Parrinello M. Canonical sampling through velocity rescaling. The Journal of Chemical Physics 126, 014101 (2007). DOI: 10.1063/1.2408420

54. Zadina JE, Hackler L, Ge LJ, Kastin AJ. A potent and selective endogenous agonist for the mu-opiate receptor. Nature 386, 499–502 (1997). DOI:10.1038/386499a0

55. Morris GM, Huey R, Lindstrom W, Sanner MF, Belew RK, et al. AutoDock4 and AutoDockTools4: Automated docking with selective receptor flexibility. Journal of Computational Chemistry 30, 2785–2791 (2009). DOI:10.1002/jcc.21256

56. Pike LJ, Han X, Chung KN, Gross RW. Lipid rafts are enriched in arachidonic acid and plasmenylethanolamine and their composition is independent of caveolin-1 expression: a quantitative electrospray ionization/mass spectrometric analysis. Biochemistry 41, 2075–2088 (2002). DOI:10.1021/bi0156557

57. Ingólfsson HI, Melo MN, van Eerden FJ, Arnarez C, Lopez CA, et al. Lipid organization of the plasma membrane. Journal of the American Chemical Society 136, 14554–14559 (2014). DOI:10.1021/ja507832e

58. Huang J, Rauscher S, Nawrocki G, Ran T, Feig M, et al. CHARMM36m: an improved force field for folded and intrinsically disordered proteins. Nature Methods 14, 71–73 (2017). DOI:10.1038/nmeth.4067

59. Bernetti M, Bussi G. Pressure control using stochastic cell rescaling. The Journal of Chemical Physics 153, 114107 (2020). DOI: 10.1063/5.0020514

60. Kabsch W, Sander C. Dictionary of protein secondary structure: Pattern recognition of hydrogen-bonded and geometrical features. Biopolymers 22, 2577–2637 (1983). DOI: 10.1002/bip.360221211

61. Grinnell SG, Uprety R, Varadi A, Subrath J, Hunkele A, Pan YX, Pasternak GW, Majumdar S. Synthesis and Characterization of Azido Aryl Analogs of IBNtxA for Radio-Photoaffinity Labeling Opioid Receptors in Cell Lines and in Mouse Brain. Cellular and Molecular Neurobiology 41, 977–993 (2021). DOI: 10.1007/s10571-020-00867-6

62. Rethi-Nagy Z, Abraham E, Udvardy K, Klement E, Darula Z, Pal M, Katona RL, Tubak V, Pali T, Kota Z, Sinka R, Udvardy A, Lipinszki Z. STABILON, a Novel Sequence Motif That Enhances the Expression and Accumulation of Intracellular and Secreted Proteins. International Journal of Molecular Sciences 23, 8168 (2022). DOI: 10.3390/ijms23158168

63. Dimmito MP, Stefanucci A, Pieretti S, Minosi P, Dvorácskó S, Tömböly C, Zengin G, Mollica A. Discovery of Orexant and Anorexant Agents with Indazole Scaffold Endowed with Peripheral Antiedema Activity. Biomolecules 9, 492 (2019). DOI: 10.3390/biom9090492

64. Borics A, Mallareddy JR, Timári I, Kövér KE, Keresztes A, Tóth G. The effect of Pro(2) modifications on the structural and pharmacological properties of endomorphin-2. Journal of Medicinal Chemistry 55, 8418–8428 (2012). DOI: 10.1021/jm300836n

65. Dvorácskó S, Keresztes A, Mollica A, Stefanucci A, Macedonio G, Pieretti S, Zádor F, Walter FR, Deli MA, Kékesi G, Bánki L, Tuboly G, Horváth G, Tömböly C. Preparation of bivalent agonists for targeting the mu opioid and cannabinoid receptors. European Journal of Medicinal Chemistry 178, 571–588 (2019). DOI: 10.1016/j.ejmech.2019.05.037

66. Borics A, Tóth G. Structural comparison of μ-opioid receptor selective peptides confirmed four parameters of bioactivity. Journal of Molecular Graphics and Modelling 28, 495–505 (2010). DOI:10.1016/j.jmgm.2009.11.006

67. Parker MS, Wong YY, Parker SL. An ion-responsive motif in the second transmembrane segment of rhodopsin-like receptors. Amino Acids 35, 1–15 (2008). DOI:10.1007/s00726-008-0637-6

68. Selent J, Sanz F, Pastor M, de Fabritiis G. Induced Effects of Sodium Ions on Dopaminergic G-Protein Coupled Receptors. PLOS Computational Biology 6, e1000884 (2010). DOI:10.1371/journal.pcbi.1000884

69. Neve KA, Cumbay MG, Thompson KR, Yang R, Buck DC, et al. Modeling and Mutational Analysis of a Putative Sodium-Binding Pocket on the Dopamine D_2_ Receptor. Molecular Pharmacology 60, 373–381 (2001). DOI:10.1124/mol.60.2.373

70. Cherezov V, Rosenbaum DM, Hanson MA, Rasmussen SGF, Thian FS, et al. High-Resolution Crystal Structure of an Engineered Human β_2_-Adrenergic G Protein–Coupled Receptor. Science 318, 1258–1265 (2007). DOI:10.1126/science.1150577

71. Palczewski K, Kumasaka T, Hori T, Behnke CA, Motoshima H, et al. Crystal Structure of Rhodopsin: A G Protein-Coupled Receptor. Science 289, 739–745 (2000). DOI:10.1126/science.289.5480.739

72. Stoddart LA, Kellam B, Briddon SJ, Hill SJ. Effect of a toggle switch mutation in TM6 of the human adenosine A_3_ receptor on G_i_ protein-dependent signalling and Gi-independent receptor internalization. British Journal of Pharmacology 171, 3827–3844 (2014). DOI:10.1111/bph.12739

73. Ma X, Hu Y, Batebi H, Heng J, Xu J, Liu X, Niu X, Li H, Hildebrand PW, Jin C, et al. Analysis of β_2_AR-G_s_ and β_2_AR-G_i_ complex formation by NMR spectroscopy. Proceedings of the National Academy of Sciences 117, 23096–23105 (2020). DOI: 10.1073/pnas.2009786117

74. Yuan S, Vogel H, Filipek S. The Role of Water and Sodium Ions in the Activation of the μ-Opioid Receptor. Angewandte Chemie 52, 10112–10115 (2013). DOI:10.1002/anie.201302244

