## Supplementary Material for "Evidencing the role of a conserved polar signaling channel in the activation mechanism of the μ-opioid receptor"

for

**Table S1.** Post-translational modifications of MOP included in the simulation systems

| Glycosylation | Phosphorylation | Lipidation / Palmitoylation |
|---|---|---|
| N9, N31, N38 (a) | S363, T370 (b) | C170$^{3.55}$ (c) |

(a) Huang P, Chen C, Xu W, Yoon SI, Unterwald EM, *et al*.. *Biochemical and Biophysical Research Communications* **365**, 82-88 (2008). DOI:10.1016/j.bbrc.2007.10.128

(b) Mann A, Illing S, Miess E, Schulz S. *British Journal of Pharmacology* **172**, 311-316 (2015). DOI:10.1111/bph.12627

(c) Zheng H, Pearsall EA, Hurst DP, Zhang Y, Chu J, *et al*. *BMC Cell Biology* **13**, 1-18 ( 2012). DOI:10.1186/1471-2121-13-6

**Table S2.** Oligonucleotide primers used in the study

| Primer name | Sequence (5'-3') | Used for |
|---|---|---|
| T7 promoter forward | TAATACGACTCACTATAGGG | Cloning MOP-1 into pJET2.1 and for sequencing |
| BGH reverse | TAGAAGGCACAGTCGAGG | |
| MOP_Y326F_fw | TGCATTGCCTTGGGTTTCACAAACAGCTGCCTG | *In vitro* mutagenesis |
| MOP_Y326F_rev | CAGGCAGCTGTTTGTGAAACCCAAGGCAATGCA | |
| MOP_N328D_fw | GCCTTGGGTTACACAGACAGCTGCCTGAACCCA | |
| MOP_N328D_rev | TGGGTTCAGGCAGCTGTCTGTGTAACCCAAGGC | |
| MOP_N328L_fw | GCCTTGGGTTACACACTCAGCTGCCTGAACCCA | |
| MOP_N328L_rev | TGGGTTCAGGCAGCTGAGTGTGTAACCCAAGGC | |
| MOP_D340N_fw | CTTTATGCGTTCCTGAATGAAAACTTCAAACGA | |
| MOP_D340N_rev | TCGTTTGAAGTTTTCATTCAGGAACGCATAAAG | |
| MOP_D340L_fw | CTTTATGCGTTCCTGCTTGAAAACTTCAAACGA | |
| MOP_D340L_rev | TCGTTTGAAGTTTTCAAGCAGGAACGCATAAAG | |
| InFusion MOP fw | CGGTACCCGGGGATCGAATTCGCCCTTGAGAGGAAGAGG | In-fusion cloning to pTRE2hyg |
| InFusion MOP rev | GCTGACTAGAGGATCGAATTCGCCCTTCAGGAAACC | |
| pTre2Hyg fw | ACGCTGTTTTGACCTCCATAG | Sequencing |
| pTre2Hyg rev | ATGAATTTTACAATAGCGAA | |

**Table S3.** $\chi^1$ Rotamer populations of aromatic amino acid side chains of EM2 during unrestrained simulations when bound to MOP derivatives.

| **TYR$^1$** | $g^+$ | $g^-$ | $t$ |
|---|---|---|---|
| **WT** | 0.00 | 0.00 | 0.98 |
| **Y326$^{7.43}$F** | 0.00 | 0.00 | 0.92 |
| **N328$^{7.45}$D** | 0.00 | 0.00 | 0.97 |
| **N328$^{7.45}$L** | 0.00 | 0.00 | 0.97 |
| **D340$^{8.47}$N** | 0.00 | 0.00 | 0.96 |
| **D340$^{8.47}$L** | 0.00 | 0.00 | 0.97 |
| **PHE$^3$** | | | |
| **WT** | 0.00 | 1.00 | 0.00 |
| **Y326$^{7.43}$F** | 0.01 | 0.18 | 0.80 |
| **N328$^{7.45}$D** | 0.01 | 0.03 | 0.95 |
| **N328$^{7.45}$L** | 0.00 | 0.93 | 0.07 |
| **D340$^{8.47}$N** | 0.00 | 0.78 | 0.21 |
| **D340$^{8.47}$L** | 0.01 | 0.93 | 0.06 |
| **PHE$^4$** | | | |
| WT | 0.00 | 1.00 | 0.00 |
| Y326$^{7.43}$F | 0.00 | 0.99 | 0.01 |
| N328$^{7.45}$D | 0.00 | 1.00 | 0.00 |
| N328$^{7.45}$L | 0.00 | 1.00 | 0.00 |
| D340$^{8.47}$N | 0.00 | 1.00 | 0.00 |
| D340$^{8.47}$L | 0.01 | 0.99 | 0.00 |

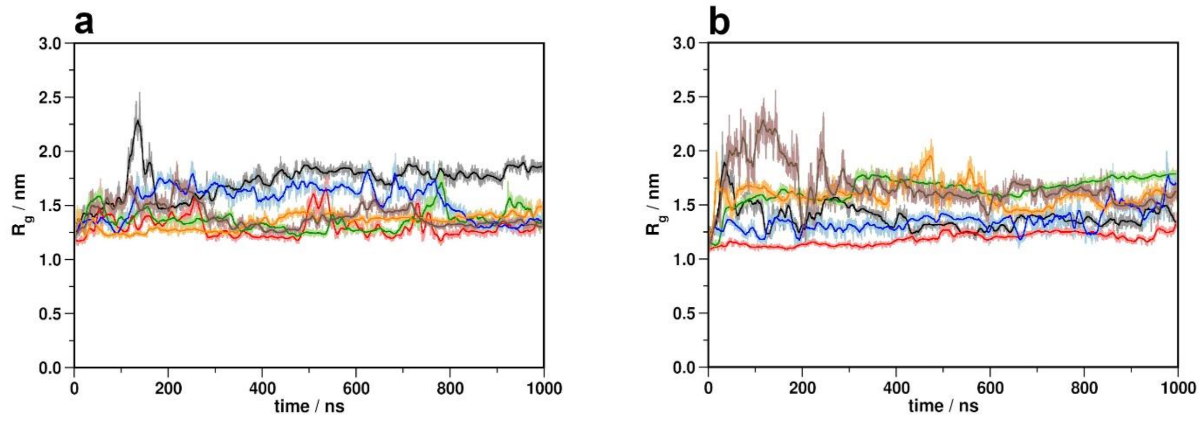


**Figure S1.** Radii of gyration of the N- (a) and C-terminal (b) domains during simulations**.** Black: wild type MOP; red: Y326$^{7.43}$F; green: N328$^{7.45}$D; blue: N328$^{7.45}$L; orange: D340$^{8.47}$N; brown: D340$^{8.47}$L.

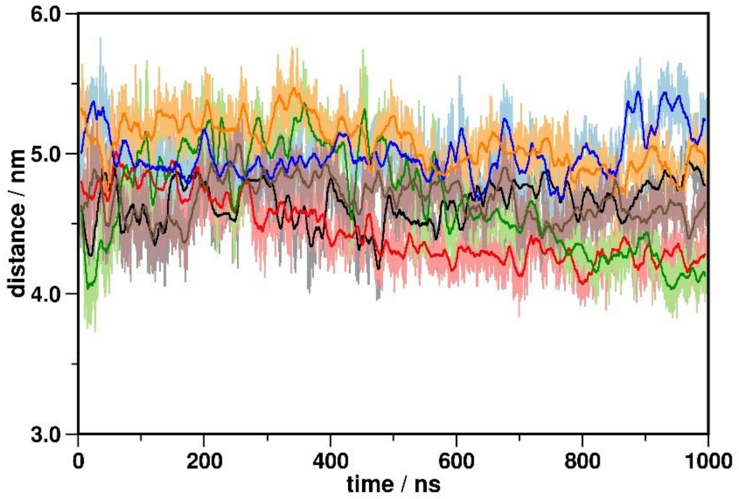


**Figure S2.** Minimum distance between the N- and C-terminal domains during simulations**.** Black: wild type MOP; red: Y326$^{7.43}$F; green: N328$^{7.45}$D; blue: N328$^{7.45}$L; orange: D340$^{8.47}$N; brown: D340$^{8.47}$L.

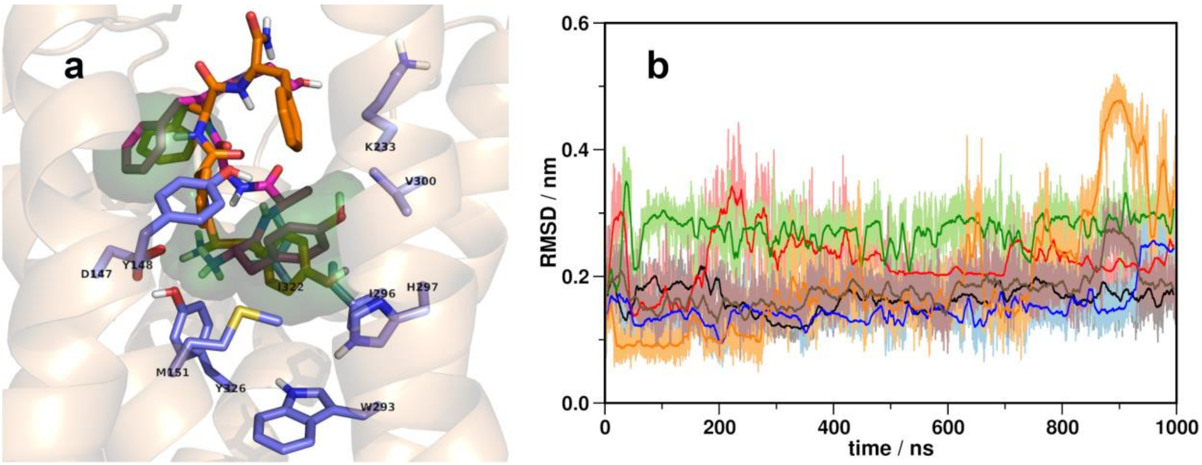


**Figure S3.** (a) Bioactive orientation of EM2 (orange) in the binding pocket of MOP, superimposed with the cryo-EM structure of DAMGO (magenta, PDB ID: 6DDE). Overlapping pharmacophore groups of the two ligands are highlighted with green. (b) Disposition of EM2 from its initial position during the course of unrestrained simulations. Black: wild type MOP; red: Y326$^{7.43}$F; green: N328$^{7.45}$D; blue: N328$^{7.45}$L; orange: D340$^{8.47}$N; brown: D340$^{8.47}$L.

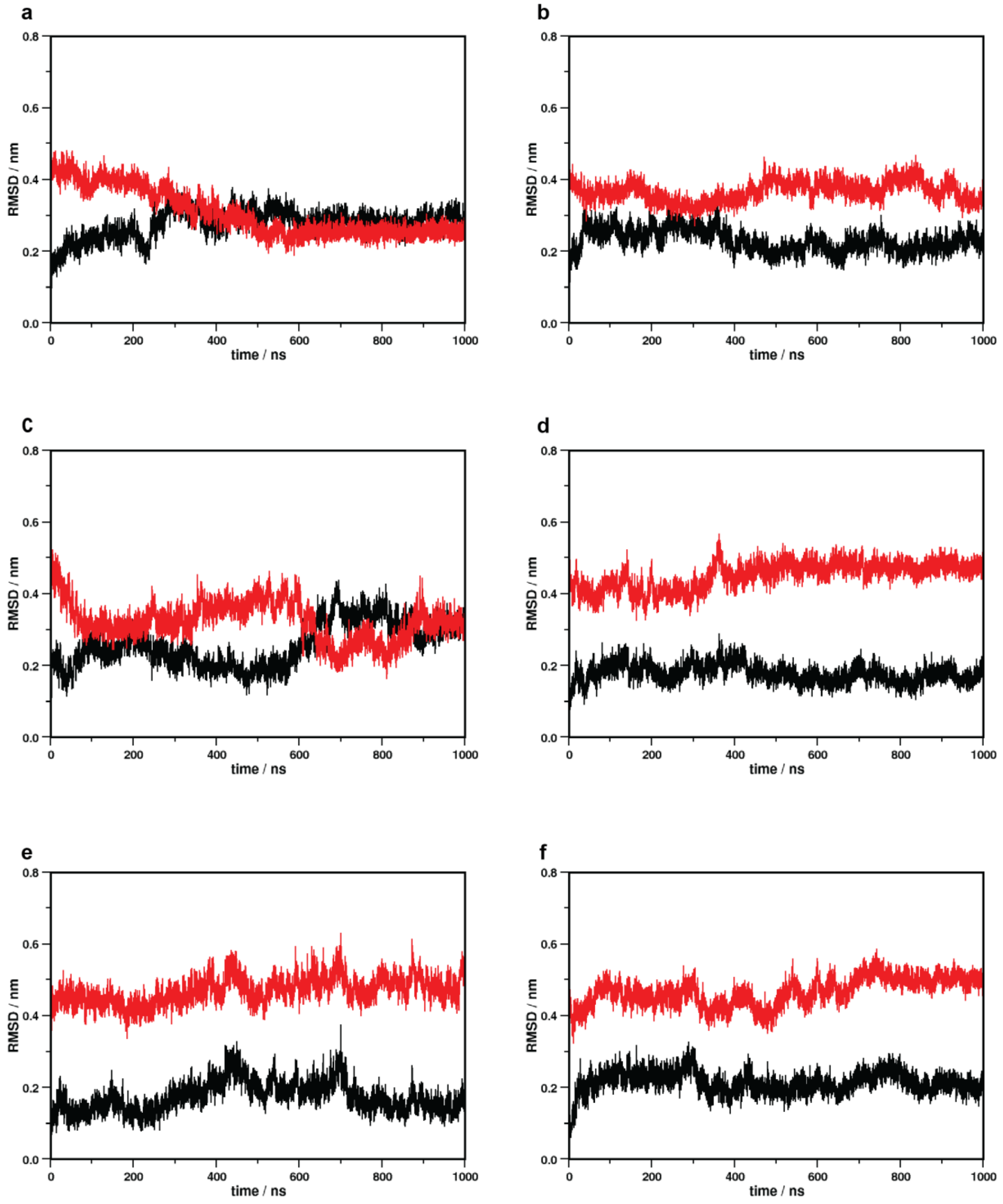


**Figure S4.** Disposition of TM6 during ligand-restrained simulations from the active (black) and the inactive (red) states. (a) wild type MOP (b) Y326$^{7.43}$F (c) N328$^{7.45}$D (d) N328$^{7.45}$L (e) D340$^{8.47}$N (f) D340$^{8.47}$L.

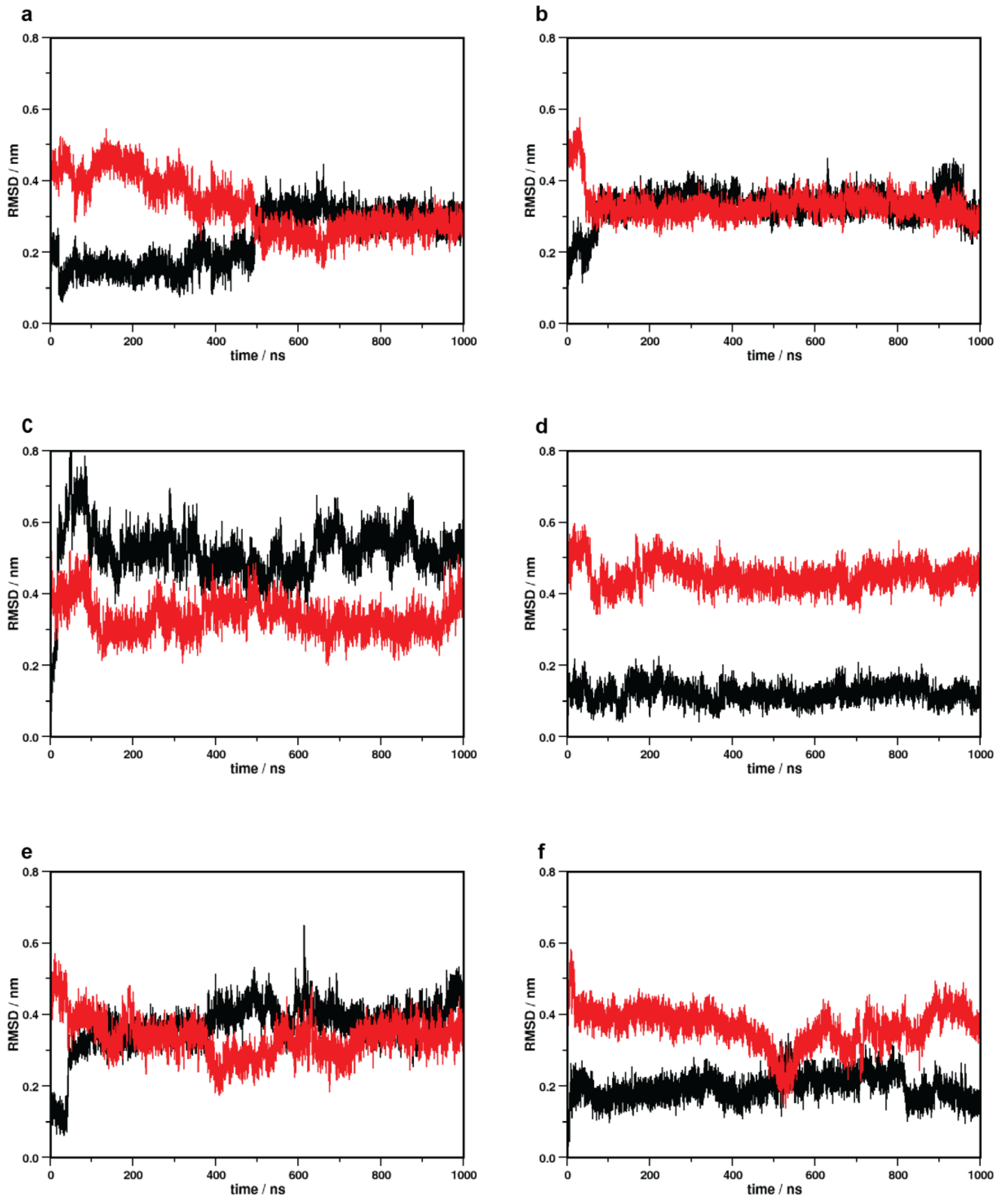


**Figure S5.** Disposition of the NPxxY motif during ligand-restrained simulations from the active (black) and the inactive (red) states. (a) wild type MOP (b) Y326$^{7.43}$F (c) N328$^{7.45}$D (d) N328$^{7.45}$L (e) D340$^{8.47}$N (f) D340$^{8.47}$L.

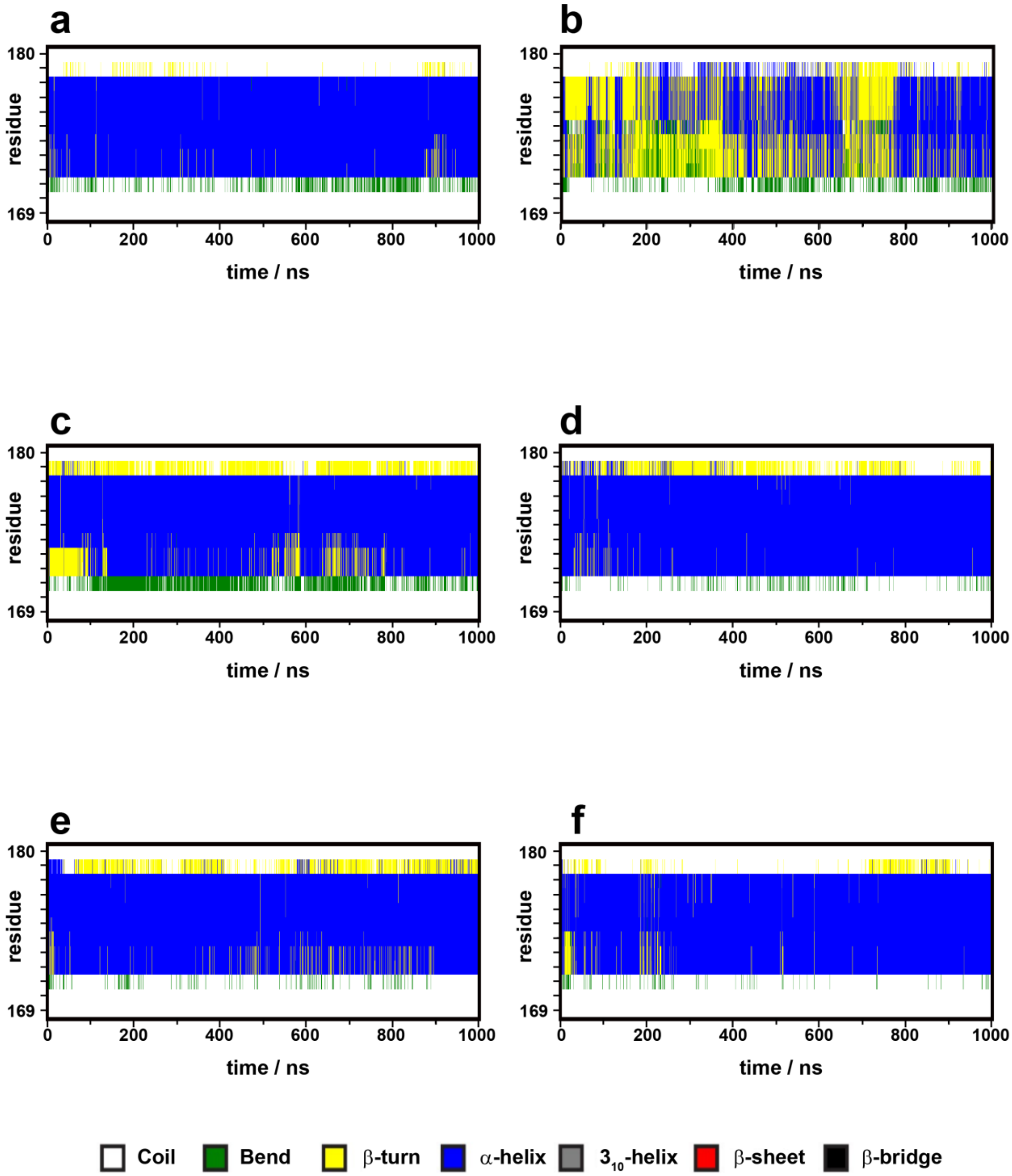


**Figure S6.** Evolution of the secondary structure of ICL2 during simulations. (a) wild type MOP (b) Y326$^{7.43}$F (c) N328$^{7.45}$D (d) N328$^{7.45}$L (e) D340$^{8.47}$N (f) D340$^{8.47}$L.

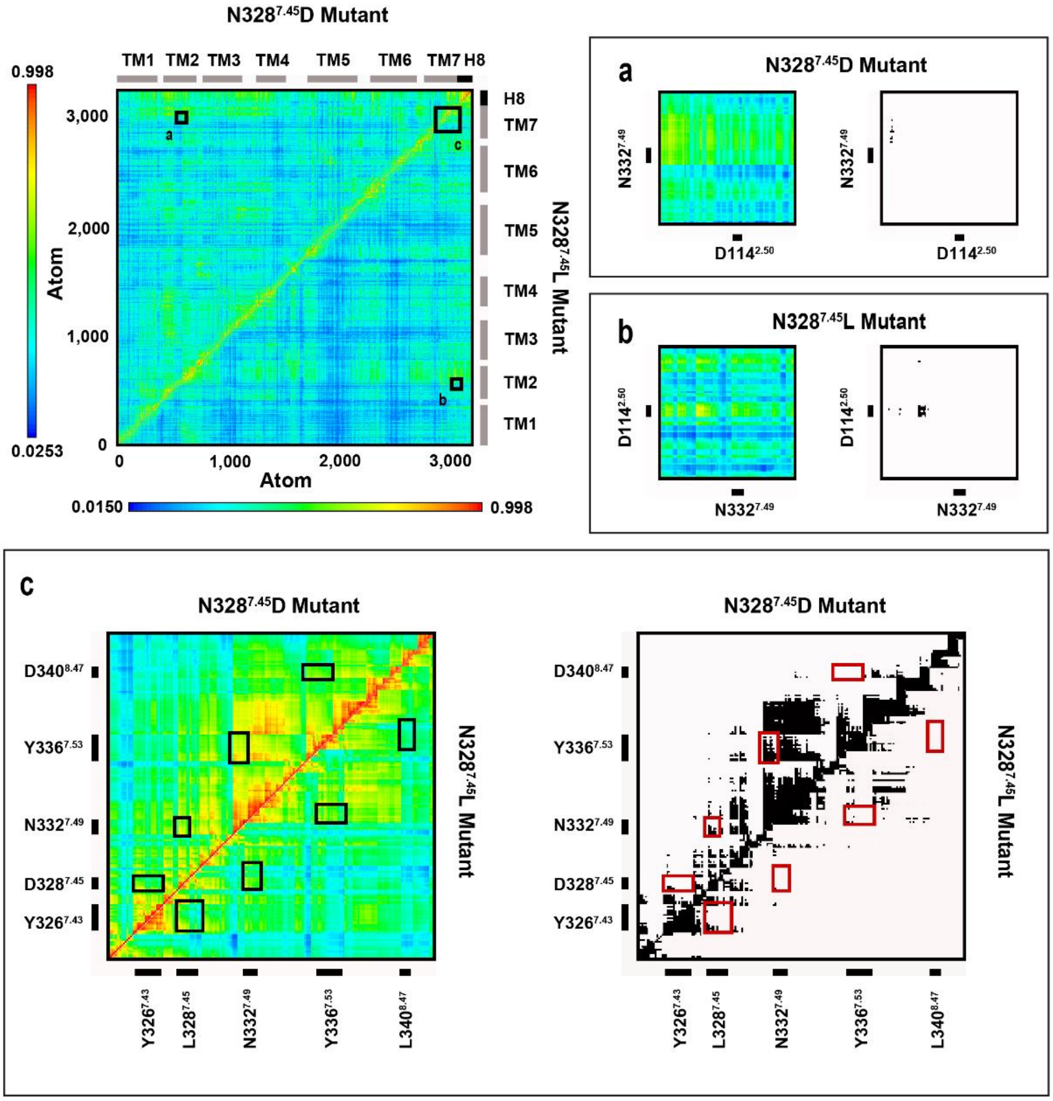


**Figure S7.** Dynamic cross-correlation matrices of the acive state, $G_i$ protein-bound N328$^{7.45}$D and N328$^{7.45}$L MOP mutants. Panels (a-c) are magnified views of regions of amino acid residues of interest. Black and white panels show correlations above the threshold of 0.63 MI.

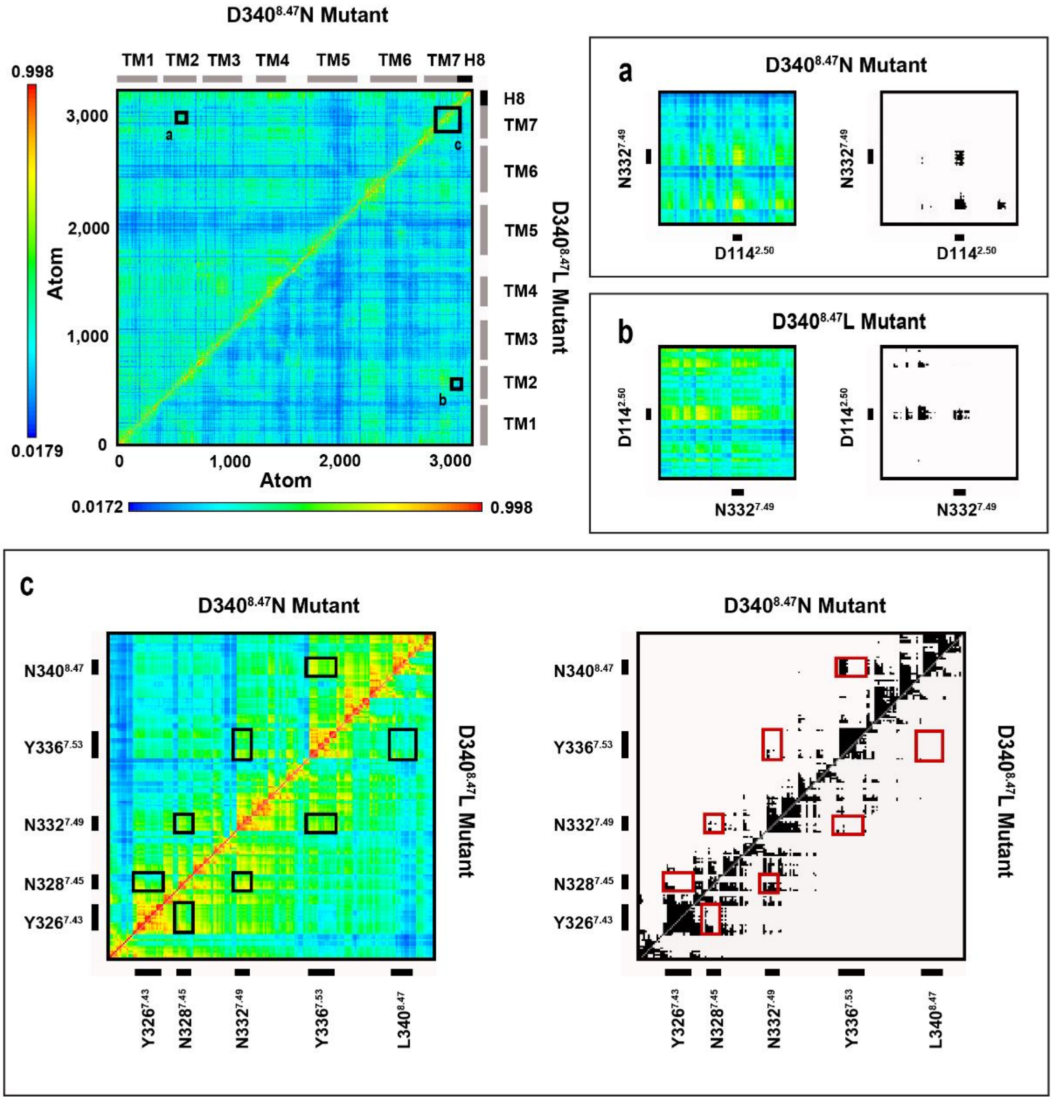


**Figure S8.** Dynamic cross-correlation matrices of the active state, $G_i$ protein-bound D340$^{8.47}$N and D340$^{8.47}$L MOP mutants. Panels (a-c) are magnified views of regions of amino acid residues of interest. Black and white panels show correlations above the threshold of 0.63 MI.

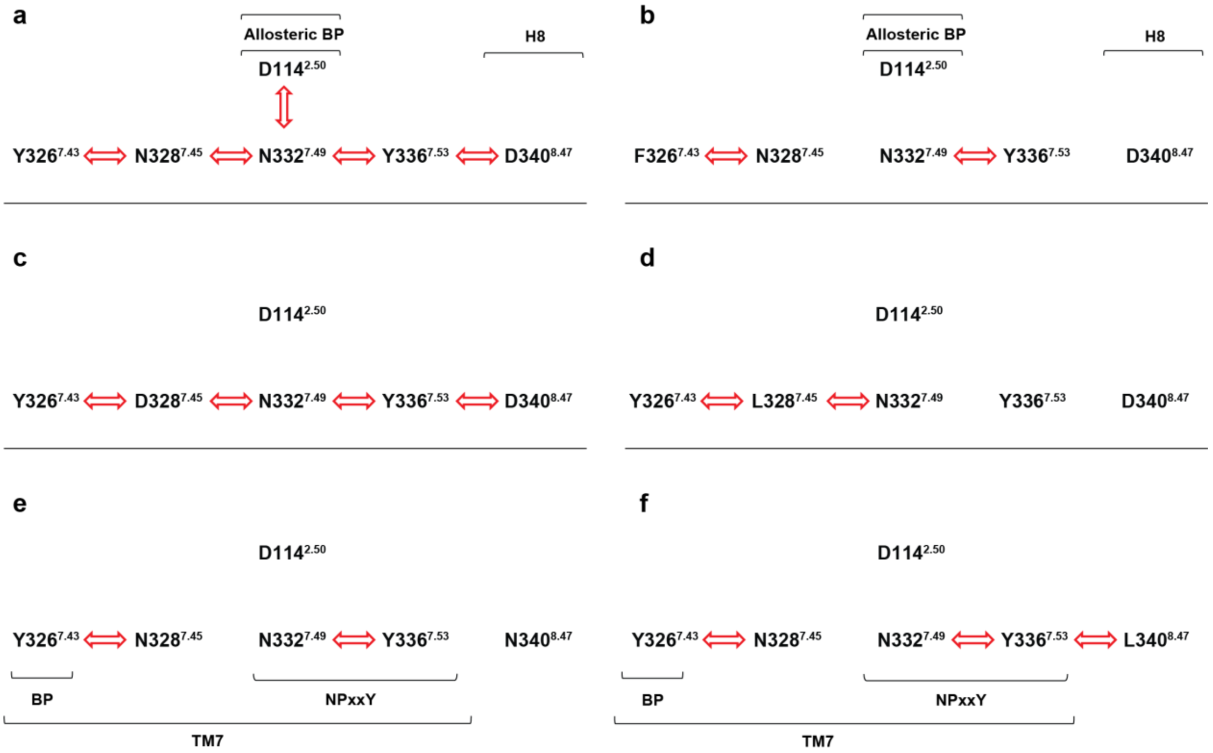


**Figure S9.** Correlated motions in the conserved polar signaling channel of active state, $G_i$ protein-bound MOP derivatives revealed by generalized cross correlation matrix analysis (ligand-restrained simulations). Red arrows indicate correlated motions of the respective amino acids in the (a) wild type MOP (b) Y326$^{7.43}$F (c) N328$^{7.45}$D (d) N328$^{7.45}$L (e) D340$^{8.47}$N (f) D340$^{8.47}$L mutants.

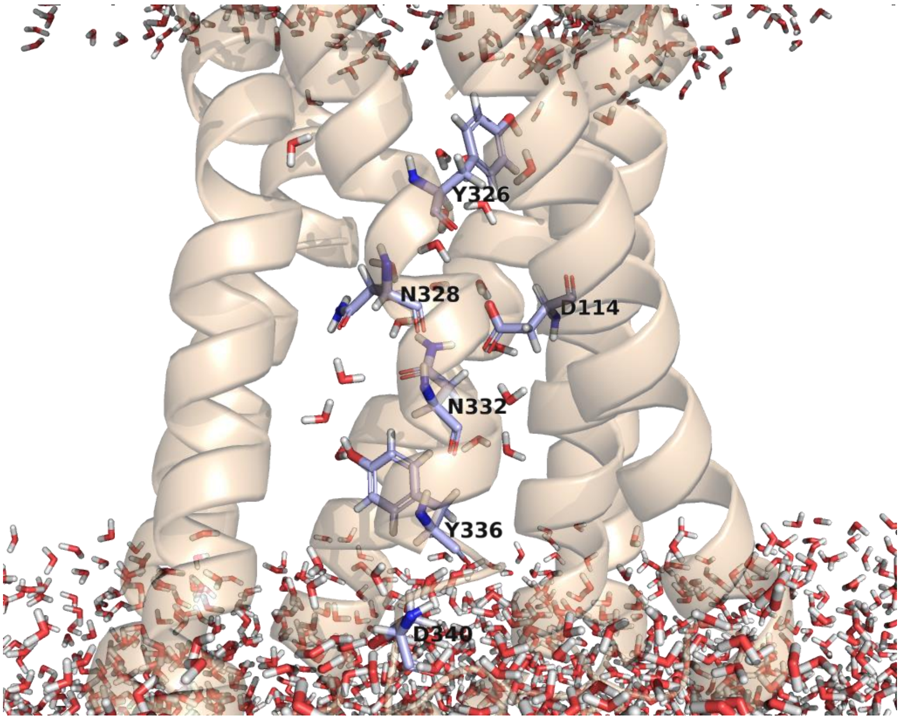


**Figure S10.** Snapshot of the transmembrane domain of the active, EM2-bound, $G_i$ protein-coupled wild type MOP. Water molecules and polar signaling channel residues in contact with water during the course of simulation are shown in stick representation.
